# Integrative Single-Cell Transcriptomics and Explainable AI for Disease Mechanism and Biomarker Discovery in Triple-Negative Breast Cancer

**DOI:** 10.64898/2026.10.23.761315

**Authors:** Tanjidul Huda, Sriporna Biswas

## Abstract

Invasion is an early step in the metastatic spread of triple-negative breast cancer (TNBC), yet cells within a single tumour or cell line differ in their capacity to invade, and it is unclear which transcriptional features distinguish cells that invade from those that do not. We re-analysed the public single-cell RNA-sequencing dataset GSE329526, in which ClonMapper-barcoded MDA-MB-231 cells were separated in a transwell assay into invasive (five samples, 6,289 cells) and non-invasive (five samples, 5,983 cells) fractions and compared with the parental population (one sample, 3,022 cells). An initial cell-level analysis combined Wilcoxon differential expression, an XGBoost classifier with a random cell split, SHapley Additive exPlanations (SHAP) and Enrichr over-representation analysis. Because cells from the same sample are not independent, we then repeated the analysis at the sample level, using differential expression on sample means (five versus five), XGBoost with cross-validation that held out whole samples, a quality-control-metric baseline model, SHAP on held-out samples and rank-based gene-set enrichment analysis (GSEA). At the sample level, 6,651 of 11,727 tested genes differed between invasive and non-invasive samples (false discovery rate [FDR] < 0.05), including the ten top-ranked cell-level genes such as *PHLDA2*, *ANTXR2*, *VIM* and *HMGA1* (FDR 0.0007–0.0022) and the epithelial-to-mesenchymal transition (EMT) regulators *TGFBR2*, *JAG1*, *SDCBP* and *WWTR1*. With whole samples held out, XGBoost distinguished invasive from non-invasive cells with an area under the ROC curve (AUC) of 0.92 (range across folds 0.89–0.95); removing ribosomal and mitochondrial genes did not change performance (AUC 0.92), whereas a model using only four quality-control metrics reached 0.63, and every held-out invasive sample received a higher mean predicted probability of invasion (0.74–0.79) than every non-invasive sample (0.20–0.28). On held-out samples, SHAP ranked inflammatory and stress-response genes highest (*CCL2*, *LGALS12*, *DUSP1*, *CXCL3*, *KYNU*, *CXCL2*), and GSEA showed higher KRAS signalling (normalised enrichment score [NES] 2.30, FDR 0.001) and E2F targets (NES 1.62, FDR 0.038) and lower interferon-α and -γ responses (NES −1.97 and −1.65; FDR 0.001 and 0.019) in invasive samples. The ribosomal genes that dominated the original SHAP ranking were absent from the held-out ranking. Invasive MDA-MB-231 cells therefore carry a transcriptional state that is reproducible across independent transwell samples and involves EMT-related, inflammatory and proliferative programmes. Limitations include a single cell line, sequencing of the invasive and non-invasive populations in separate libraries and the absence of external validation; the candidate genes reported here are hypotheses that require confirmation in independent experiments and patient material.

## INTRODUCTION

Breast cancer is the most commonly diagnosed cancer in women worldwide [1]. Triple-negative breast cancer (TNBC), defined by the absence of oestrogen and progesterone receptors and of HER2 amplification, accounts for a minority of cases but a disproportionate share of early recurrence and death, and it lacks the targeted treatment options available for other subtypes [2,3]. Most of this mortality is caused by metastasis, which begins when tumour cells leave the primary site and invade the surrounding tissue. Understanding which cells acquire invasive behaviour, and which transcriptional programmes accompany it, is therefore central to understanding TNBC progression.

TNBC is biologically heterogeneous. Expression profiling divides it into several molecular subtypes, and widely used TNBC cell lines represent different subtypes, MDA-MB-231 being classified as mesenchymal stem-like [4]. Heterogeneity also exists within a single tumour or cell population: not every cell invades, and cells that share a genetic background can differ in their capacity to migrate. Whether such differences are reflected in a reproducible transcriptional state, and whether that state can be learned and interpreted by machine learning, is a distinct question from the classification of tumour subtypes.

In this study we re-analysed single-cell transcriptomes of invasive and non-invasive MDA-MB-231 cells from a lineage-tracing transwell experiment. The study makes four contributions: (i) a description of transcriptional differences between invasive and non-invasive cells, first at the cell level and then at the level of independent samples; (ii) an XGBoost classifier of invasion status evaluated with cross-validation that holds out whole samples, together with a quality-control-metric baseline that measures technical separability; (iii) interpretation of the classifier with SHAP values computed on held-out samples; and (iv) rank-based pathway analysis and a comparison of the resulting candidate genes with published TNBC evidence. Rather than proposing validated biomarkers, the aim is to establish which differences are reproducible across samples and to show how the validation design changes the conclusions of an explainable-AI analysis.

## BACKGROUND

Single-cell RNA sequencing (scRNA-seq) measures the expression of thousands of genes in individual cells and has revealed subclonal structure and aggressive cell states within TNBC tumours [5] as well as the cellular composition of breast cancers more broadly [6]. Expressed DNA barcodes add lineage information to these measurements: with ClonMapper, every cell carries a heritable barcode that is captured in its transcriptome, so that clonal identity and gene expression can be read from the same cell [7]. Combined with a functional assay such as the transwell invasion assay, in which cells that migrate through a porous membrane towards a serum gradient are separated from those that remain, this design links a cell’s behaviour, its clone and its transcriptome [8].

Machine-learning classifiers are increasingly applied to scRNA-seq data to find genes that distinguish cell states. Gradient-boosted trees such as XGBoost handle many correlated features and non-linear interactions [9], and SHAP values attribute each prediction to individual features, which makes such models easier to interpret [10,11]. A feature that a model relies on is not necessarily a biological marker of the class it predicts, however, and two methodological issues determine how far such results can be trusted.

First, cells from the same sample are not independent replicates. Statistical tests that treat thousands of cells as independent observations produce many false discoveries, and aggregating cells into one profile per sample (pseudobulk analysis) is recommended for comparisons between conditions [12,13]. Second, when related observations appear in both training and test data, or when class labels coincide with technical batches, machine-learning performance is overestimated, a problem known as data leakage [14]. In single-cell studies both issues arise when cells are split at random, because every sample then contributes cells to training and testing, and a model can succeed by recognising sample-specific or library-specific signals rather than the biological state of interest.

## LITERATURE REVIEW

Single-cell studies of TNBC have shown that tumours contain transcriptionally distinct subpopulations, some of which are linked to aggressive behaviour and treatment resistance [5], and breast cancer atlases have mapped malignant cell states across subtypes [6]. Molecular classification of TNBC into subtypes with distinct biology and preferred cell-line models [4] provides the context for studying a single model such as MDA-MB-231. Lineage barcoding has made it possible to follow clones through selection and treatment and to ask whether particular clones are enriched under a given pressure [7]; the dataset analysed here applied this approach to invasion [8].

Explainable machine learning has been used to prioritise genes that drive model predictions, with tree-based models and SHAP values a common combination [9–11]. Methodological work has, however, shown that single-cell differential expression is prone to false discoveries when cells are treated as replicates [12,13] and that leakage between training and test data is a widespread cause of irreproducible machine-learning results [14]. Together, these studies imply that explainable models of single-cell data should be evaluated on samples they were not trained on, and that the features they use should be compared with sample-level statistics before biological conclusions are drawn.

The present study differs from earlier work in its question and design. It addresses invasion within a single TNBC cell line rather than differences between tumours or subtypes; it uses a public lineage-traced dataset with five independent samples per group; and it reports a cell-level analysis alongside a sample-aware re-analysis, so that the effect of the validation design on performance, feature importance and pathway results can be seen directly. We do not claim that this combination is unique; rather, it provides a transparent and reproducible baseline for further studies of invasive cell states.

## METHODOLOGY

### Dataset and study design

The processed AnnData file GSE329526_MB231.h5ad was obtained from the Gene Expression Omnibus (series GSE329526; Illumina NovaSeq 6000; public since 6 May 2026) [8]. According to the GEO record, ClonMapper-barcoded MDA-MB-231 cells were seeded in transwell chambers with a fetal bovine serum gradient (0.5% to 20%), and cells that had migrated through the membrane after 24 hours were collected as invasive, cells that had not as non-invasive; a parental population was profiled separately. The parental, invasive and non-invasive populations were each sequenced as a separate gene-expression library with sample multiplexing. No associated publication was listed at the time of analysis. The object contained 15,294 cells and 36,601 genes from 11 samples (Table 1). In the object, the field *label* assigns samples C3, C5, C6, C7 and C8 to Invasive, B3, B5, B6, B7 and B8 to NonInvasive, and PT to Control; we refer to the PT sample as the parental population. Lineage groups (High-Invasive, Low-Invasive, Non-Invasive and TopLineage; field *AbunInvLineage*), 380 lineage barcodes (*collapsedLineages*) and Leiden clusters were provided by the data generators; their definitions are not documented in the GEO record.

**Table 1.**
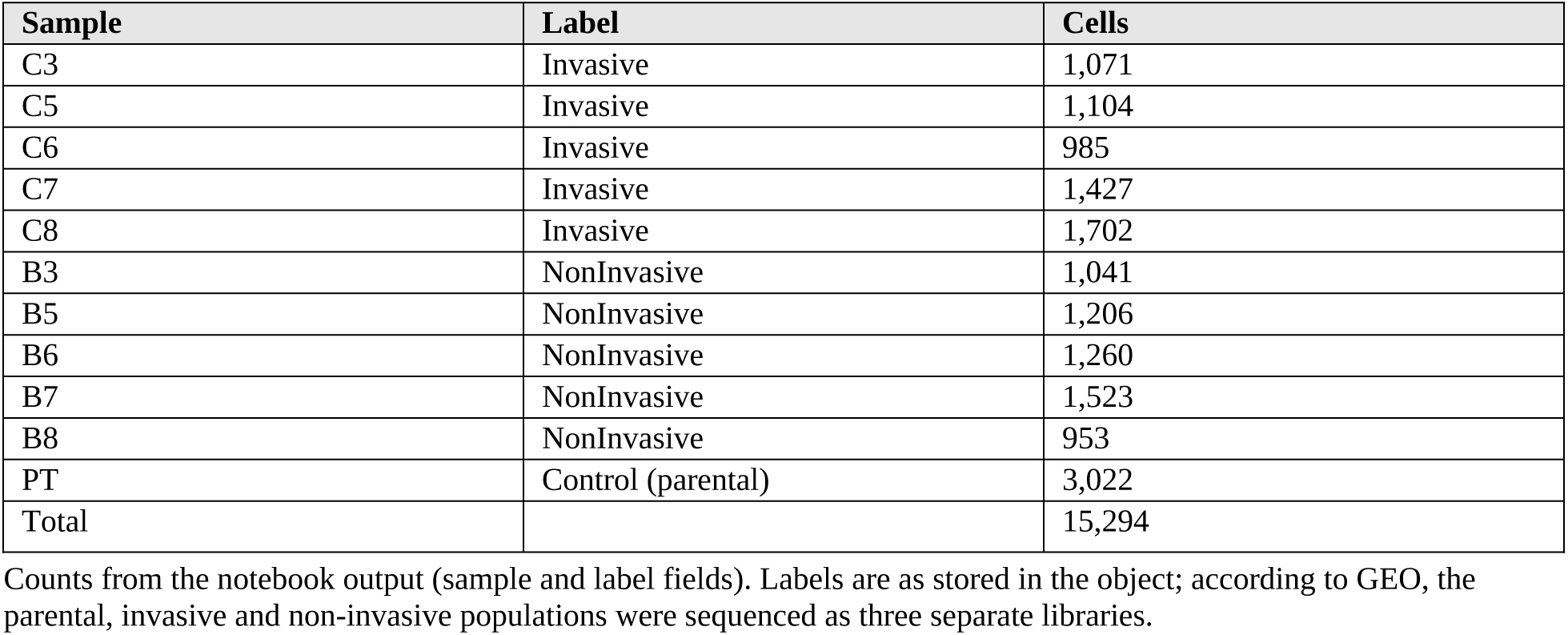
Samples and cell numbers in GSE329526_MB231.h5ad.

### Pre-processing

Quality-control metrics, outlier flags, normalisation, highly variable gene selection (2,491 genes), principal-component analysis, neighbourhood graph, UMAP and Leiden clustering were already present in the deposited object, and we did not repeat or modify them. No cell was flagged as an outlier, indicating that flagged cells had already been removed. The object contains no raw counts: *adata.X* is identical to the layer *log1p_norm*, with values between 0 and 1.16 and mean values of about 1.0 for highly expressed genes (for example 1.03 for *VIM* and 1.04 for *GAPDH*). This range is compressed relative to log1p-transformed counts per 10,000, so expression differences are reported on the deposited scale and are not interpreted as log_2_ fold changes. Methods requiring raw counts (for example DESeq2) could not be applied.

### Cell-level differential expression

Genes were ranked between Invasive and NonInvasive cells with the Wilcoxon rank-sum test in Scanpy (*rank_genes_groups*, *method=’wilcoxon’*, Invasive versus reference NonInvasive) [15], with Benjamini–Hochberg adjustment across all 36,601 genes [16]. Each cell was treated as an observation. The same function was used to compare each lineage group with all remaining cells (one-versus-rest, grouped by *AbunInvLineage*). Scanpy computes log fold changes on the assumption that the input is log1p-transformed; given the uncertainty about the scale of *adata.X*, reported fold changes are used only to indicate direction.

### Classification with XGBoost

An XGBoost classifier (*multi:softprob*, 50 trees, maximum depth 4, other parameters at their defaults, random seed 42) [9] was trained to predict the three labels (Control, Invasive, NonInvasive) from all 36,601 genes. Cells were split at random into training (80%; 12,235 cells) and test (20%; 3,059 cells) sets with *train_test_split* (random state 42, not stratified). No hyper-parameter tuning, cross-validation or class weighting was performed. Performance on the test cells was summarised by accuracy, per-class precision, recall and F1-score, the confusion matrix, and one-versus-rest ROC curves with areas under the curve (AUC). Because the split was made at the cell level, every sample contributed cells to both the training and the test set.

### SHAP interpretation

SHAP values were computed with the TreeExplainer of the SHAP package [10,11] for 100 test cells drawn at random without a fixed seed. Gene importance was summarised as the mean absolute SHAP value across the 100 cells and the three classes. Summary (beeswarm) plots were produced for the Control class (class index 0) only.

### Functional enrichment

Over-representation analysis was performed with Enrichr [17] through GSEApy 1.2.1 [18] against GO Biological Process 2023 [19] and KEGG 2021 Human [20] gene-set libraries. Three input lists were analysed: (i) the 200 genes with the highest Wilcoxon scores for Invasive versus NonInvasive cells, all of which were higher in invasive cells; (ii) the 50 genes with the largest mean absolute SHAP values; and (iii) the 20 genes with the highest scores for the High-Invasive lineage group versus all other cells. Enrichr’s default background was used; no experiment-specific background of detected genes was supplied. Enrichr adjusted p-values (Benjamini– Hochberg) below 0.05 were considered significant.

### Protein–protein interaction network

A protein–protein interaction network was built in STRING [21], and node degree was used to rank hub genes. The gene list submitted, the minimum interaction score, the evidence channels and the software used to compute degree were not recorded in the notebook; the degree values for the ten top-ranked genes were entered manually for plotting. Because these details are missing, the network result is reported as descriptive only.

### Literature comparison

Selected candidate genes were compared with published studies of TNBC or basal-like breast cancer identified by targeted searches. Only studies whose bibliographic details could be verified are cited, and the type of evidence (experimental or observational) is recorded.

### Sample-level differential expression

To avoid treating cells as independent replicates [12,13], we computed for each of the ten invasive and non-invasive samples the mean expression of every gene detected in at least 10% of cells in either group (11,727 genes), and compared the five invasive with the five non-invasive sample means with Welch’s t-test and Benjamini–Hochberg correction.

### Sample-aware classification

XGBoost (binary logistic objective, 50 trees, maximum depth 4, histogram tree method, seed 42) was trained to distinguish invasive from non-invasive cells using the 2,491 highly variable genes. Performance was estimated with five-fold cross-validation grouped by well number, so that each fold held out one invasive and one non-invasive sample (C3 with B3, C5 with B5, and so on) and no sample contributed cells to both training and test data. We report the pooled AUC and balanced accuracy over all held-out cells, the AUC of each fold, and the mean predicted probability of invasion for each held-out sample. The same procedure was applied (i) after removing ribosomal-protein and mitochondrial genes (2,485 genes) and (ii) to four quality-control metrics alone (total counts, detected genes, mitochondrial and ribosomal percentages), as a baseline for technical separability. For comparison, the binary model was also evaluated with a random stratified 80/20 cell split. The parental class was excluded because it consists of a single sample.

### SHAP on held-out samples

For the first cross-validation fold (samples B7 and C7 held out), SHAP values of the HVG model were computed with TreeExplainer for 1,000 held-out cells selected with a fixed seed, and genes were ranked by mean absolute SHAP value.

### Rank-based enrichment

Genes were ranked by the sample-level t statistic and analysed with GSEA prerank in GSEApy (1,000 permutations, gene-set size 15–500, seed 42) against the MSigDB Hallmark 2020, GO Biological Process 2023 and KEGG 2021 libraries. Gene sets with FDR < 0.25 are reported, following the GSEA convention. Enrichr over-representation analysis of the sample-level significant genes, with the tested genes as background, was also performed.

### Software

The original analysis was run in Google Colab (Python 3.12) with Scanpy, XGBoost, SHAP, scikit-learn 1.6.1, GSEApy 1.2.1, pandas 2.3.3, NumPy 2.0.2 and matplotlib 3.10.0; the Scanpy, XGBoost and SHAP versions of that run were not recorded. The sample-aware re-analysis was run locally with Python 3.12.15, Scanpy 1.12.1, NumPy 2.5.4, pandas 3.0.6, scikit-learn 1.9.1, XGBoost 3.4.1, SHAP 0.54.1 and GSEApy 1.2.1. Re-running the original three-class model in this environment gave an accuracy of 0.965 (original 0.966).

## RESULTS AND DISCUSSION

### Dataset composition

The 15,294 cells came from five invasive samples (6,289 cells; 985–1,702 per sample), five non-invasive samples (5,983 cells; 953–1,523 per sample) and one parental sample (3,022 cells) (Table 1). The data generators’ lineage annotation assigned 4,348 cells to the High-Invasive group, 5,798 to Low-Invasive, 1,095 to Non-Invasive and 4,053 to TopLineage; the TopLineage group corresponds to the single most abundant barcode (4,053 cells). Leiden clustering at resolution 0.04 separated the cells into two clusters (Supplementary Figure S1). The overall workflow is shown in Figure 1.

**Figure 1.**
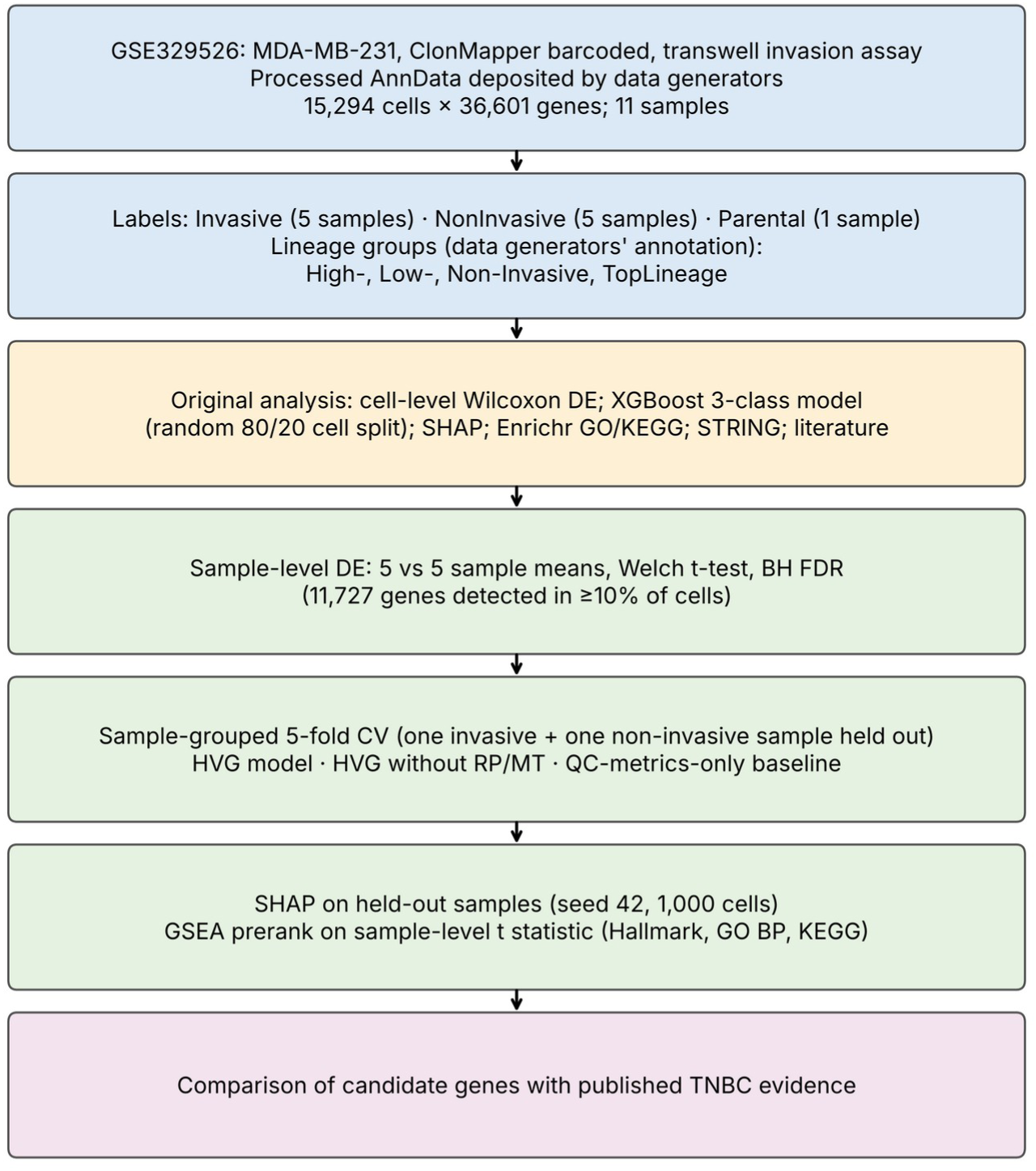
Workflow of the study. The original cell-level analysis (orange) was followed by a sample-aware re-analysis (green). DE, differential expression; CV, cross-validation; HVG, highly variable genes; RP/MT, ribosomal-protein and mitochondrial genes; BH, Benjamini–Hochberg; GSEA, gene-set enrichment analysis.

### Cell-level differential expression

Of 36,601 genes, 8,546 had an adjusted p-value below 0.05 at the cell level (2,332 higher and 6,214 lower in invasive cells), and 12 genes had adjusted p-values that were reported as exactly zero. Fold changes were mostly small: 294 of the significant genes had an absolute log fold change above 1 (Figure 2), and the 200 top-ranked genes had log fold changes between 0.02 and 1.22. The highest-ranked genes in invasive cells were *PHLDA2*, *ANTXR2*, *VIM*, *HMGA1*, *CALM1*, *MANCR*, *ARHGDIB*, *FOXP1*, *ATP5MC2*, *RPSA* (Table 2). Their expression across the three labels is shown in Figure 3. With roughly 6,000 cells per group, very small differences reach extreme significance, and these p-values do not account for the five-sample structure of each group; the number of genes that differ consistently between invasive and non-invasive samples was therefore tested at the sample level.

**Figure 2.**
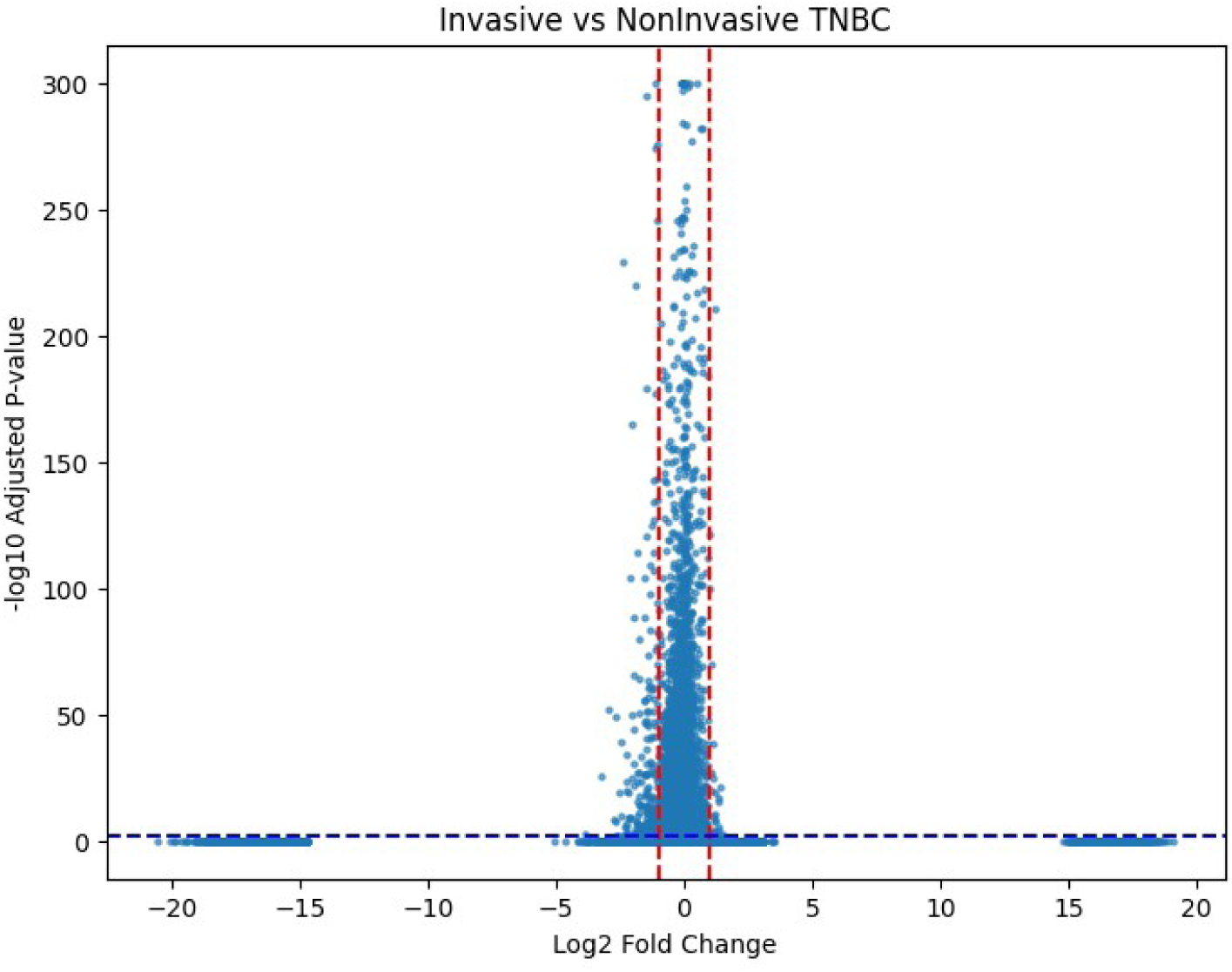
Volcano plot of the cell-level Wilcoxon comparison of invasive versus non-invasive cells (36,601 genes). Red dashed lines, log fold change ±1; blue dashed line, adjusted p = 0.01. Adjusted p-values below 10^−300^ are plotted at 300. The points near ±15–20 are genes detected in only one group. The x-axis label was written as log_2_ fold change in the original plot; values are on the deposited normalised scale.

**Figure 3.**
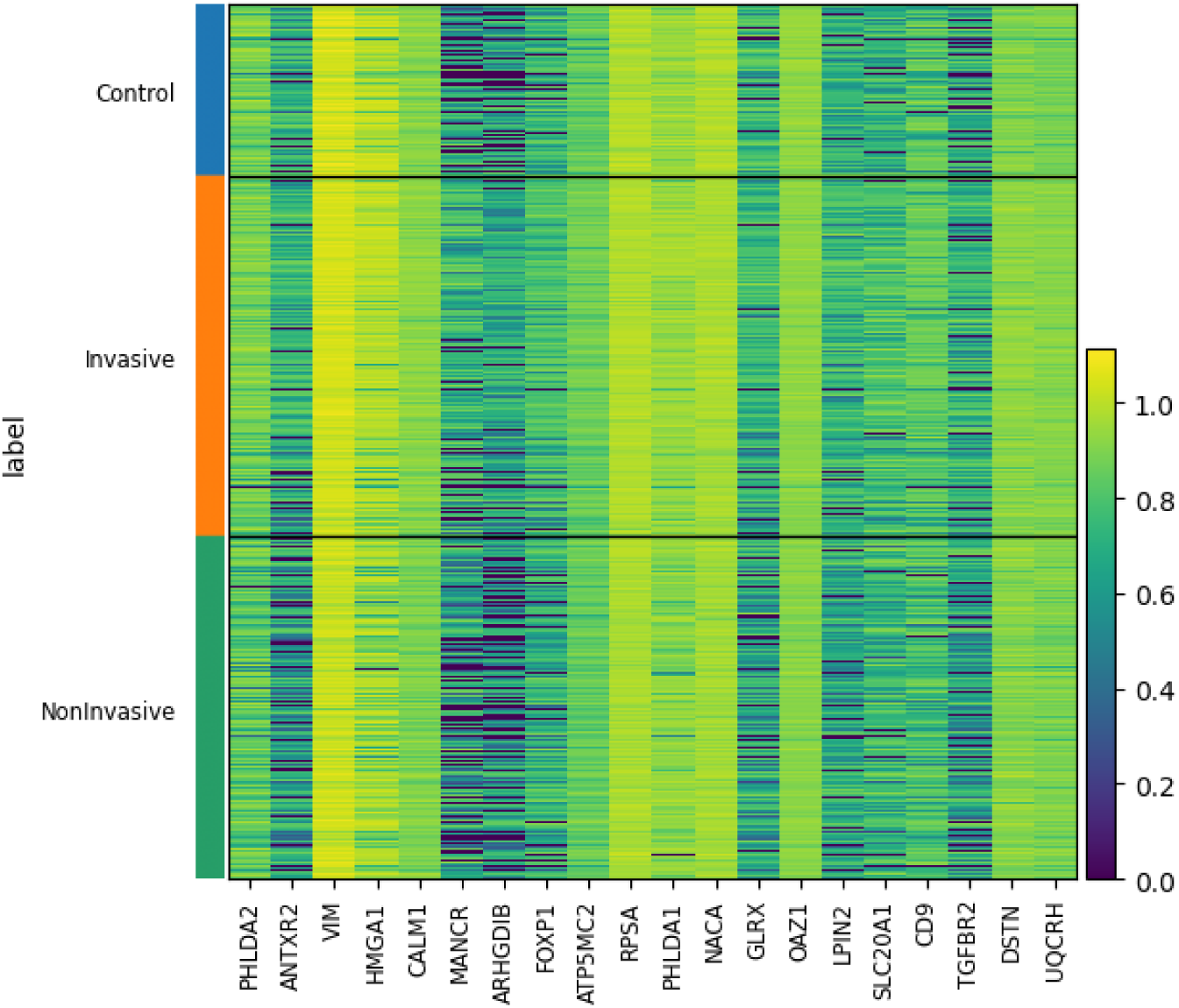
Expression (adata.X) of the 20 top-ranked genes from Table 2 in individual cells, grouped by label (Control, parental; Invasive; NonInvasive).

**Table 2.** Top 20 genes ranked by Wilcoxon score, Invasive versus NonInvasive cells (cell-level test)

| Gene | Wilcoxon score | Log fold change | Adjusted p |
| --- | --- | --- | --- |
| *PHLDA2* | 39.3 | 0.192 | $<1 \times 10^{-300}$ |
| *ANTXR2* | 38.0 | 0.513 | $<1 \times 10^{-300}$ |
| *VIM* | 37.8 | 0.049 | $<1 \times 10^{-300}$ |
| *HMGA1* | 37.2 | 0.142 | $2.05 \times 10^{-299}$ |
| *CALM1* | 36.2 | 0.061 | $5.25 \times 10^{-284}$ |
| *MANCR* | 36.2 | 0.664 | $6.71 \times 10^{-283}$ |
| *ARHGDIB* | 36.1 | 0.704 | $2.22 \times 10^{-282}$ |
| *FOXP1* | 35.8 | 0.321 | $1.33 \times 10^{-277}$ |
| *ATP5MC2* | 34.7 | 0.072 | $4.51 \times 10^{-260}$ |
| *RPSA* | 34.3 | 0.031 | $1.99 \times 10^{-254}$ |
| *PHLDA1* | 34.0 | 0.065 | $3.07 \times 10^{-250}$ |
| *NACA* | 33.8 | 0.036 | $2.86 \times 10^{-247}$ |
| *GLRX* | 33.0 | 0.357 | $2.65 \times 10^{-236}$ |
| *OAZ1* | 32.9 | 0.036 | $1.67 \times 10^{-234}$ |
| *LPIN2* | 32.8 | 0.276 | $2.43 \times 10^{-232}$ |
| *SLC20A1* | 32.3 | 0.217 | $2.38 \times 10^{-226}$ |
| *CD9* | 32.3 | 0.176 | $4.50 \times 10^{-226}$ |
| *TGFBR2* | 32.3 | 0.348 | $7.08 \times 10^{-226}$ |
| *DSTN* | 32.2 | 0.051 | $1.03 \times 10^{-224}$ |
| *UQCRH* | 32.1 | 0.053 | $1.02 \times 10^{-223}$ |

Source: Invasive\_vs\_NonInvasive\_DEG.csv. Positive fold changes indicate higher expression in invasive cells. Fold changes were computed by Scanpy from adata.X, which is on a compressed normalised scale (see Methods). Adjusted p-values treat cells as independent observations.

### Sample-level differential expression

Comparing the five invasive with the five non-invasive sample means, 6,651 of 11,727 tested genes had an FDR below 0.05 (1,313 higher and 5,338 lower in invasive samples). The differences were consistent across samples but small on the deposited scale: 252 genes were higher by more than 0.05 units and 772 lower by more than 0.05 units. All ten top-ranked cell-level genes remained significant (FDR 0.0007–0.0022; Table 3), as did the EMT-related genes *TGFBR2*, *JAG1*, *SDCBP*, *WWTR1* and *MDK* (FDR ≤ 0.012). Among the genes with the largest differences, *AGR2*, *ARHGDIB*, *MANCR*, *KRT81*, *SOX4* and *FOXA2* were higher in invasive samples, and *DSP*, *SAA1*, *SAA2*, *TNFSF10*, *NEDD9*, *THBS1* and *IL32* were lower. The large number of significant genes (all results in Supplementary Table S4) indicates a broad, consistent shift between the two populations; it may also include library-level differences, because all invasive samples were sequenced in one library and all non-invasive samples in another.

**Table 3.** Selected genes in the sample-level comparison (five invasive versus five non-invasive samples)

| Gene | Mean, invasive | Mean, non-invasive | Difference | FDR | Group |
| --- | --- | --- | --- | --- | --- |
| *PHLDA2* | 0.877 | 0.802 | +0.075 | 0.001 | Top-10 cell level |
| *ANTXR2* | 0.684 | 0.524 | +0.161 | 0.002 | Top-10 cell level |
| *VIM* | 1.035 | 1.014 | +0.021 | 0.001 | Top-10 cell level |
| *HMGA1* | 0.974 | 0.915 | +0.059 | 0.002 | Top-10 cell level |
| *MANCR* | 0.646 | 0.455 | +0.190 | 0.002 | Top-10 cell level |
| *ARHGDIB* | 0.611 | 0.417 | +0.194 | 0.002 | Top-10 cell level |
| *FOXP1* | 0.748 | 0.637 | +0.111 | 0.002 | Top-10 cell level |
| *TGFBR2* | 0.669 | 0.557 | +0.112 | 0.003 | EMT regulation (GO) |
| *JAG1* | 0.627 | 0.559 | +0.067 | 0.002 | EMT regulation (GO) |
| *SDCBP* | 0.692 | 0.642 | +0.049 | 0.001 | EMT regulation (GO) |
| *WWTR1* | 0.632 | 0.582 | +0.049 | 0.012 | EMT regulation (GO) |
| *MDK* | 0.660 | 0.588 | +0.072 | 0.010 | EMT regulation (GO) |
| *AGR2* | 0.671 | 0.465 | +0.206 | 0.002 | Largest increase |
| *KRT81* | 0.824 | 0.636 | +0.188 | $7.27 \times 10^{-4}$ | Largest increase |
| *SOX4* | 0.462 | 0.301 | +0.161 | 0.007 | Largest increase |
| *FOXA2* | 0.506 | 0.337 | +0.169 | 0.002 | Largest increase |
| *DSP* | 0.292 | 0.567 | -0.275 | 0.004 | Largest decrease |
| *SAA1* | 0.329 | 0.580 | -0.250 | 0.002 | Largest decrease |
| *TNFSF10* | 0.263 | 0.504 | -0.241 | 0.001 | Largest decrease |
| *NEDD9* | 0.150 | 0.373 | -0.222 | $8.82 \times 10^{-4}$ | Largest decrease |
| *THBS1* | 0.250 | 0.454 | -0.204 | 0.001 | Largest decrease |

Source: pseudobulk\_DE\_invasive\_vs\_noninvasive.csv. Means of per-sample average expression on the deposited normalised scale; Welch t-test, Benjamini–Hochberg FDR across 11,727 genes.

### Classification of invasion status

In the original analysis, on the 3,059 test cells, the XGBoost classifier reached an accuracy of 0.966, compared with 0.414 for always predicting the most frequent class (Table 4). Most errors were confusions between invasive and non-invasive cells (48 invasive cells predicted as non-invasive and 41 non-invasive cells predicted as invasive), whereas 569 of 582 parental cells were classified correctly (Figure 4A). One-versus-rest AUCs were 1.000 for the parental class, 0.994 for invasive and 0.996 for non-invasive cells (Figure 4B).

**Figure 4.**
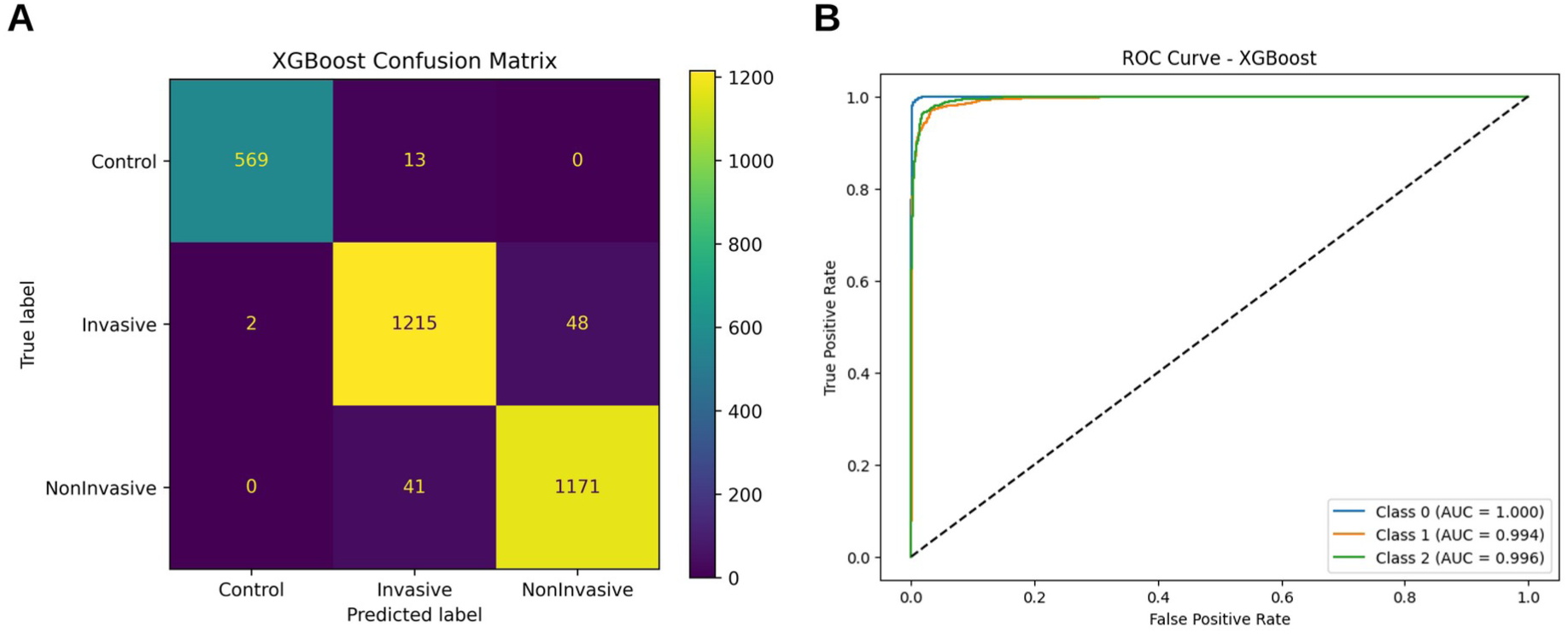
XGBoost classification of 3,059 held-out cells after a random cell-level split. (A) Confusion matrix. (B) One-versus-rest ROC curves; class 0, Control (parental); class 1, Invasive; class 2, NonInvasive. The dashed line shows chance performance.

**Table 4.**
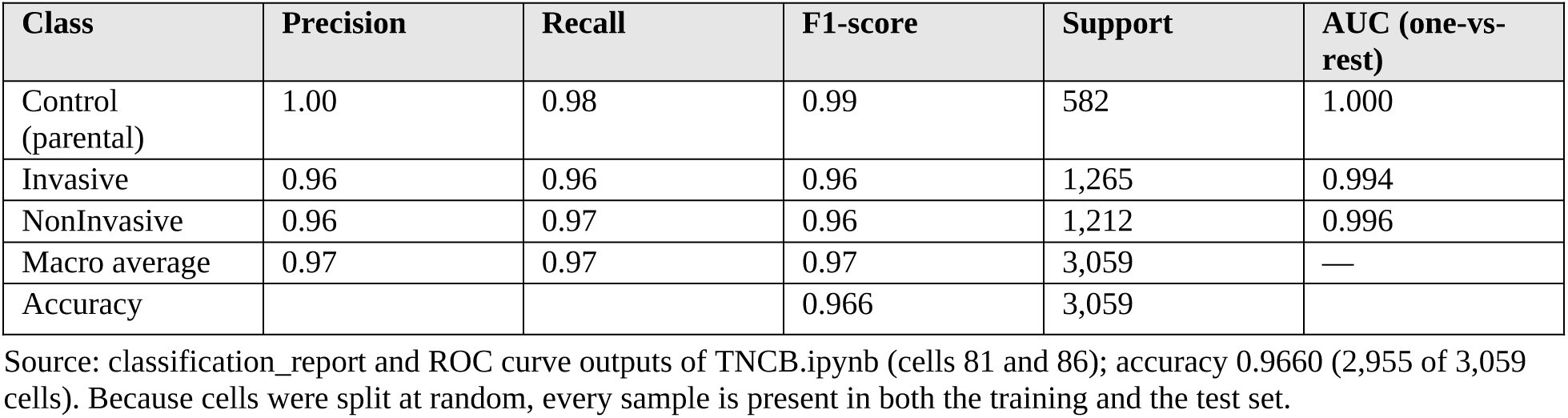
XGBoost performance on the random 20% cell-level test set.

| Class | Precision | Recall | F1-score | Support | AUC (one-vs-rest) |
| --- | --- | --- | --- | --- | --- |
| Control (parental) | 1.00 | 0.98 | 0.99 | 582 | 1.000 |
| Invasive | 0.96 | 0.96 | 0.96 | 1,265 | 0.994 |
| NonInvasive | 0.96 | 0.97 | 0.96 | 1,212 | 0.996 |
| Macro average | 0.97 | 0.97 | 0.97 | 3,059 | — |
| Accuracy |  |  | 0.966 | 3,059 |  |

Source: classification\_report and ROC curve outputs of TNCB.ipynb (cells 81 and 86); accuracy 0.9660 (2,955 of 3,059 cells). Because cells were split at random, every sample is present in both the training and the test set.

These estimates should be interpreted with caution. Test cells came from the same samples, and therefore the same sequencing libraries, as the training cells, and each label corresponds to one library. The model can therefore succeed by learning sample- or library-specific features, and its accuracy is an upper bound rather than an estimate of performance on new samples. In particular, the parental class consists of one sample, so its near-perfect separation cannot be distinguished from a library effect.

### Classification with whole samples held out

When whole samples were held out, the HVG model distinguished invasive from non-invasive cells with a pooled AUC of 0.92 and a balanced accuracy of 0.84; fold AUCs ranged from 0.89 to 0.95 (Table 5; Figure 5A). Each held-out sample was classified correctly on average: the mean predicted probability of invasion was 0.74–0.79 for the five invasive samples and 0.20–0.28 for the five non-invasive samples, and 82–89% of cells in invasive samples and 12–21% of cells in non-invasive samples were called invasive (Figure 5B; Supplementary Table S1). With a random cell-level split the same model reached an AUC of 0.93, so sample-level leakage inflated performance only slightly. Removing ribosomal-protein and mitochondrial genes left performance unchanged (AUC 0.92). A model trained on four quality-control metrics alone reached an AUC of 0.63, reflecting a modest technical difference between the populations (median 39,000–43,000 counts per cell in invasive samples versus 49,000–51,000 in non-invasive samples); this explains only a small part of the separation achieved with gene expression.

**Figure 5.**
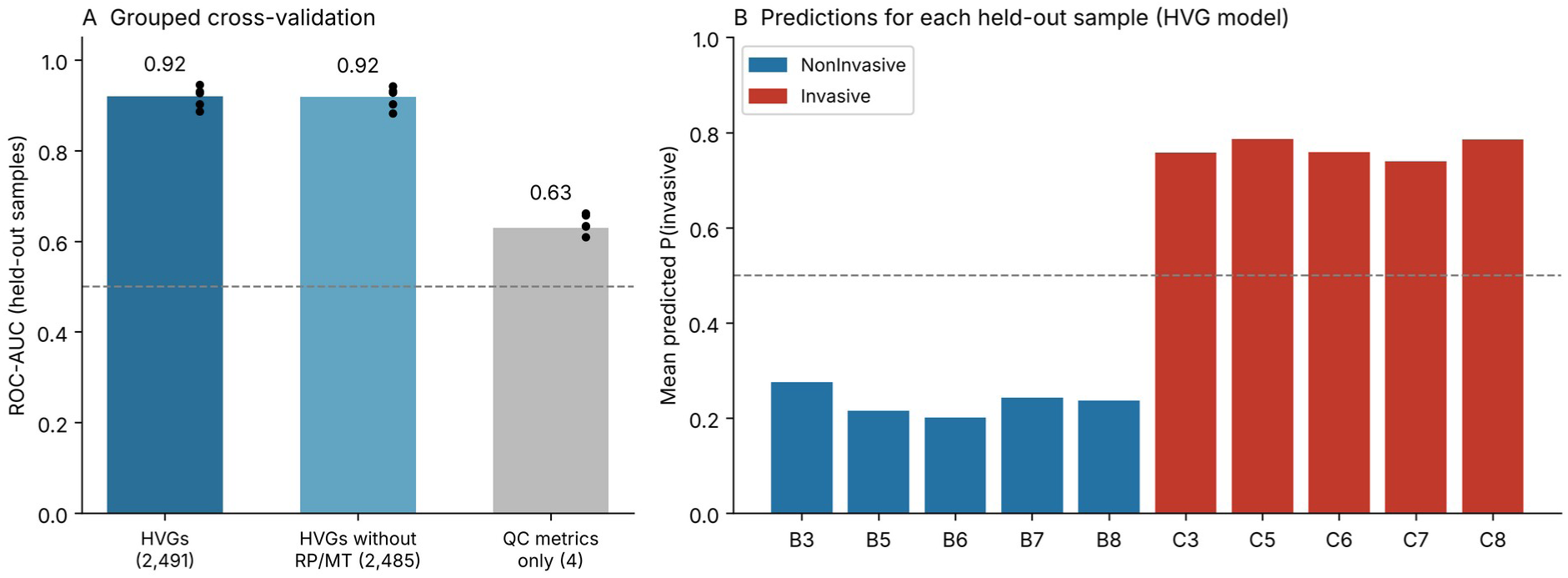
Classification with whole samples held out. (A) Pooled ROC-AUC for three feature sets; dots show the AUC of each of the five folds; the dashed line marks chance. (B) Mean predicted probability of invasion for every held-out sample (HVG model).

**Table 5.** Invasive versus non-invasive classification with sample-aware validation.

| Model | Features | Pooled AUC | Fold AUC range | Balanced accuracy |
| --- | --- | --- | --- | --- |
| XGBoost, grouped CV | 2,491 HVGs | 0.919 | 0.887–0.945 | 0.840 |
| XGBoost, grouped CV | 2,485 HVGs without RP/MT | 0.919 | 0.882–0.942 | 0.840 |
| XGBoost, grouped CV | 4 QC metrics | 0.630 | 0.610–0.662 | 0.589 |
| XGBoost, random cell split | 2,491 HVGs | 0.931 | — | 0.855 |

Source: corrected\_results.json. Grouped CV: five folds, each holding out one invasive and one non-invasive sample with the same well number. RP/MT, ribosomal-protein and mitochondrial genes; QC metrics, total counts, detected genes, mitochondrial and ribosomal percentages.

### SHAP interpretation of the original model

The 20 genes with the largest mean absolute SHAP values are listed in Table 6. Sixteen of them were ribosomal-protein or mitochondrially encoded genes (for example *RPS21*, *RPS28*, *RPL37* and *MT-ND3*), and the remaining four were *NACA*, *COX7A2*, *ATP5F1E* and *DUSP1*; across the top 50, 28 genes were ribosomal or mitochondrial. Most of these genes showed very small differences between invasive and non-invasive cells in the cell-level test, and several ranked among the genes most strongly lower in invasive cells (Table 6). Mean expression of the top 20 SHAP genes differed by only a few percent between labels (Supplementary Figure S2). The beeswarm plot in Figure 6 shows SHAP values for the parental class only; feature-value colouring was not rendered in the original plot, so the direction of each gene’s effect cannot be read from it.

**Figure 6.**
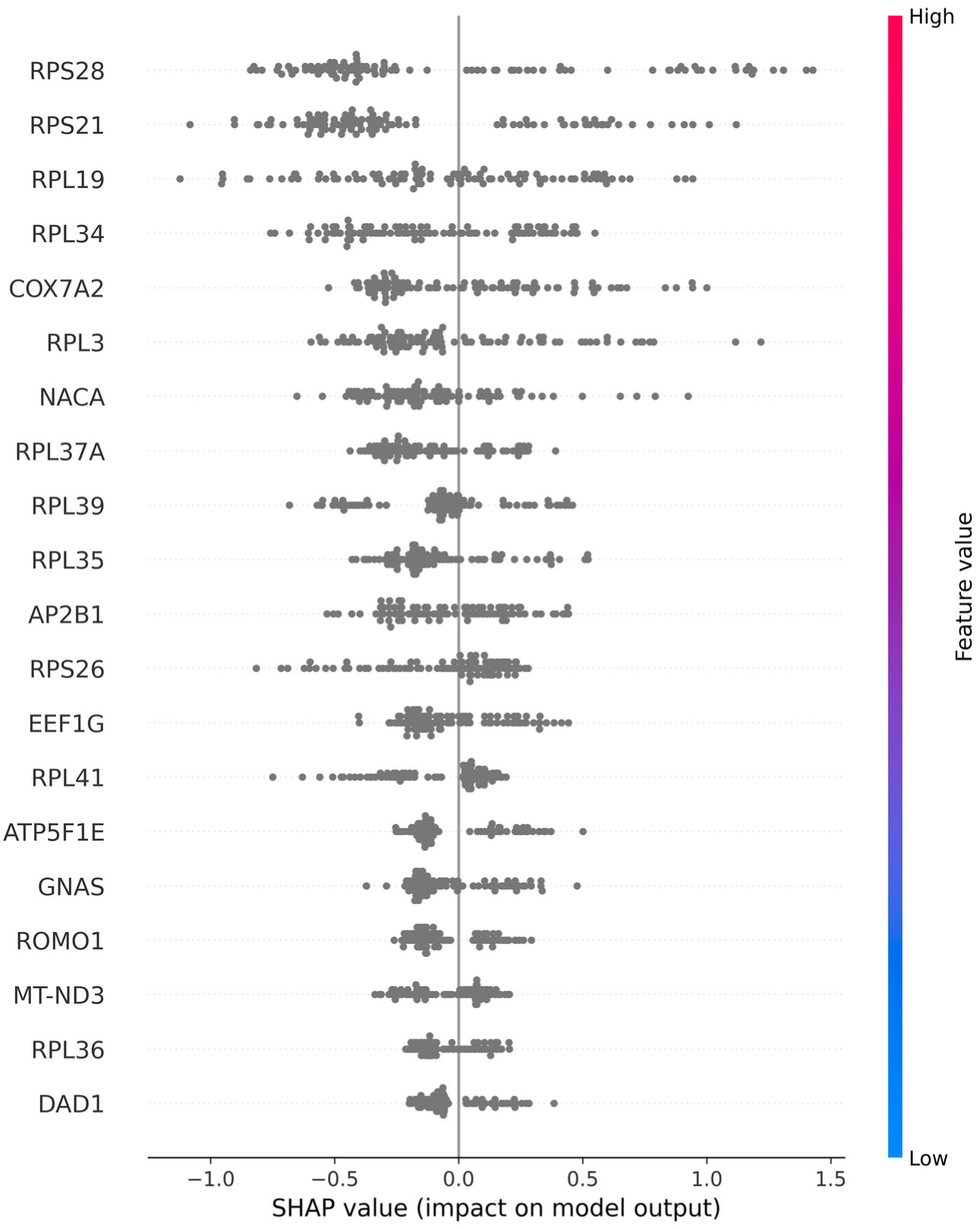
SHAP summary plot for the Control (parental) class: SHAP values of the 20 most important genes for this class in 100 randomly selected test cells. Each point is one cell; positive values push the prediction towards the parental class. Feature-value colours were not rendered (all points grey).

**Table 6.**
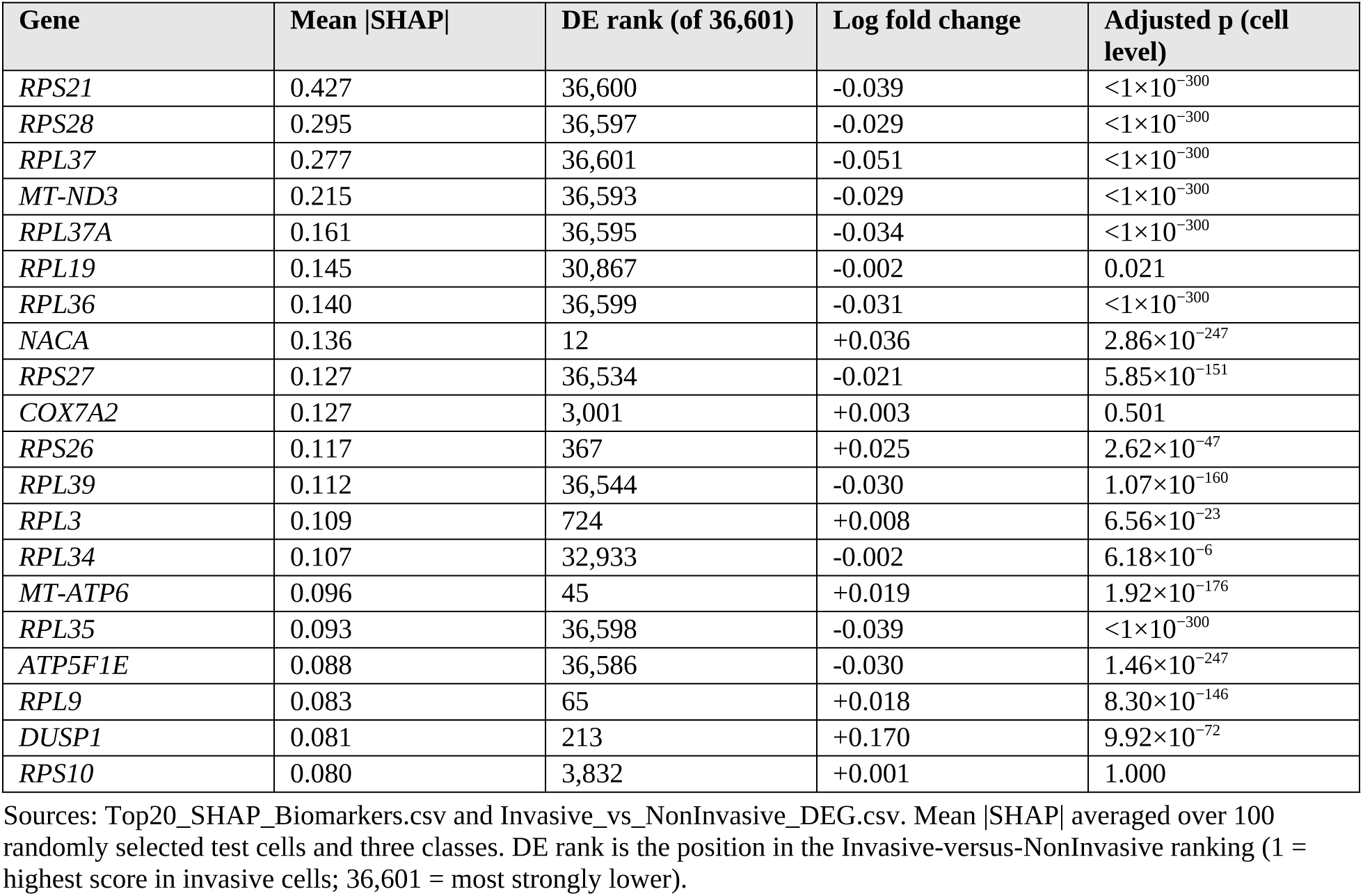
Top 20 genes by mean absolute SHAP value and their cell-level differential expression.

Ribosomal-protein and mitochondrial transcript fractions are standard technical quality metrics in scRNA-seq because they vary with cell state, cell integrity and library preparation [22]. Their dominance among the most important features is consistent with the model relying partly on cell-quality differences. The sample-aware analysis addressed this directly: removing these genes did not reduce performance, and they were absent from the held-out SHAP ranking.

### SHAP interpretation on held-out samples

For the model evaluated on held-out samples B7 and C7, none of the 20 genes with the largest mean absolute SHAP values was a ribosomal or mitochondrial gene (Table 7; Figure 7; the top 50 are listed in Supplementary Table S3). The ranking was led by *CCL2*, *LGALS12*, *DUSP1*, *MTRNR2L8*, *PRSS2*, *CXCL3*, *KYNU* and *CXCL2*, and included *NFKBIZ*, *JUN*, *DDIT4*, *EPHB6*, *NEDD9* and *AGR2*. Higher expression of *CXCL3*, *CXCL2*, *KYNU*, *NFKBIZ*, *POU2F2*, *AGR2* and *EPHB6* pushed predictions towards the invasive class, whereas *CCL2*, *LGALS12*, *NEDD9*, *LSR* and *CCDC3* were lower in invasive samples. All of these genes except *EPHB6* (FDR 0.31) also differed at the sample level. Several of them (*CXCL2*, *CXCL3*, *NFKBIZ*, *JUN*, *DUSP1*) are NF-κB targets or immediate-early genes.

**Figure 7.**
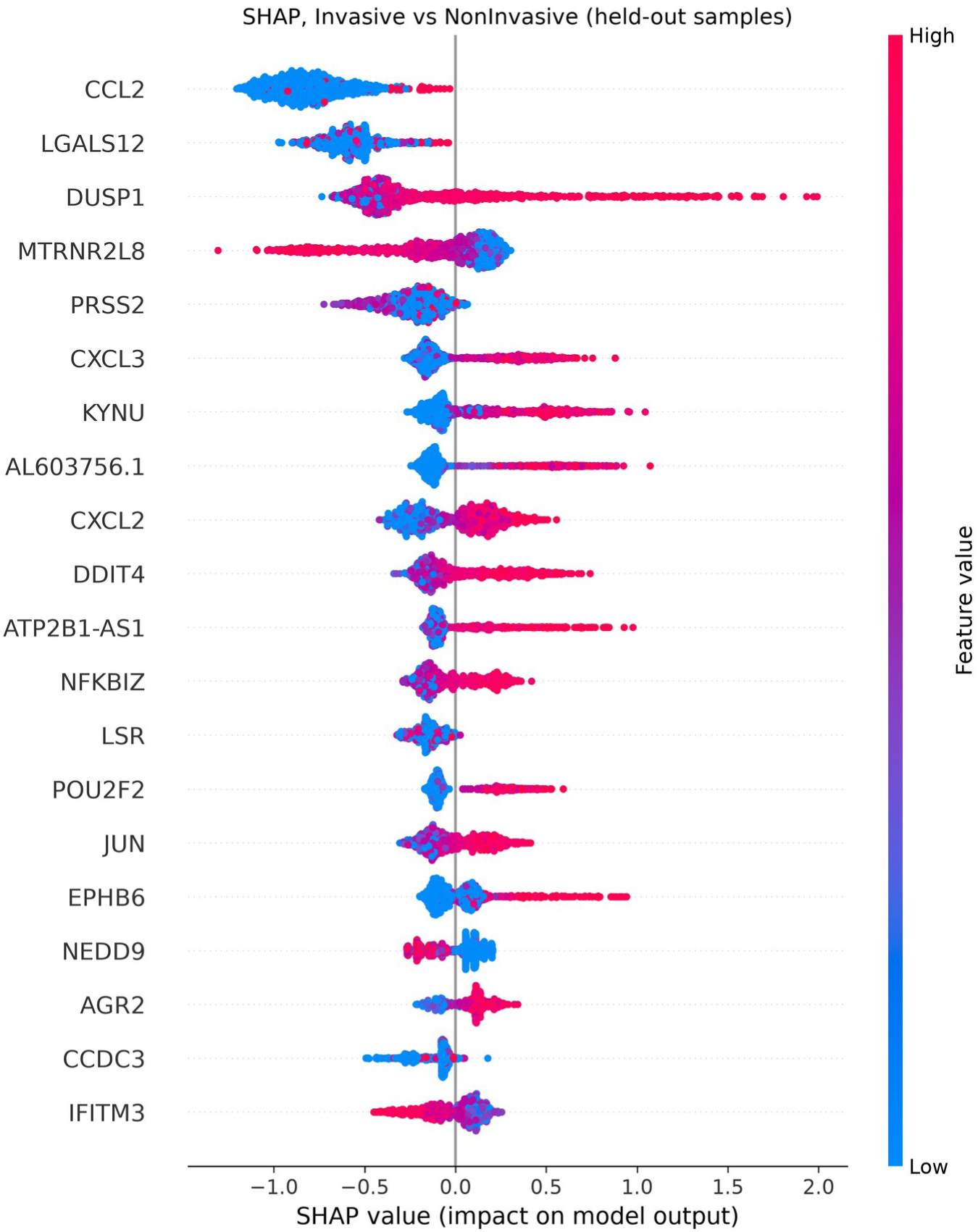
SHAP summary plot for the Invasive-versus-NonInvasive model on 1,000 cells from held-out samples B7 and C7. Each point is one cell; positive SHAP values push the prediction towards the invasive class; colour shows the gene’s expression in that cell (red, high; blue, low).

**Table 7.** Top 20 genes by mean absolute SHAP value on held-out samples (Invasive versus NonInvasive model)

| Gene | Mean \|SHAP\| | Sample-level difference | FDR |
| --- | --- | --- | --- |
| *CCL2* | 0.803 | -0.064 | $4.62 \times 10^{-4}$ |
| *LGALS12* | 0.576 | -0.205 | $8.78 \times 10^{-4}$ |
| *DUSP1* | 0.423 | +0.054 | 0.034 |
| *MTRNR2L8* | 0.240 | -0.143 | $1.43 \times 10^{-4}$ |
| *PRSS2* | 0.237 | -0.105 | 0.016 |
| *CXCL3* | 0.215 | +0.127 | 0.004 |
| *KYNU* | 0.214 | +0.170 | 0.002 |
| *AL603756.1* | 0.207 | +0.088 | 0.001 |
| *CXCL2* | 0.189 | +0.080 | 0.030 |
| *DDIT4* | 0.184 | +0.014 | 0.027 |
| *ATP2B1-AS1* | 0.156 | +0.069 | 0.038 |
| *NFKBIZ* | 0.151 | +0.123 | 0.001 |
| *LSR* | 0.150 | -0.141 | 0.001 |
| *POU2F2* | 0.145 | +0.090 | 0.002 |
| *JUN* | 0.136 | +0.043 | 0.025 |
| *EPHB6* | 0.126 | +0.035 | 0.315 |
| *NEDD9* | 0.124 | -0.222 | $8.82 \times 10^{-4}$ |
| *AGR2* | 0.121 | +0.206 | 0.002 |
| *CCDC3* | 0.120 | -0.043 | $4.49 \times 10^{-4}$ |
| *IFITM3* | 0.120 | -0.033 | 0.005 |

Sources: shap\_rank\_invasive\_vs\_noninvasive\_heldout.csv and pseudobulk\_DE\_invasive\_vs\_noninvasive.csv. 1,000 cells from held-out samples B7 and C7. Difference, invasive minus non-invasive sample mean on the deposited scale.

### Functional enrichment

In the original over-representation analysis, the 200 genes ranked highest in invasive cells were enriched for ten GO Biological Process terms and eleven KEGG pathways at an adjusted p-value below 0.05 (Table 8). Besides translation-related terms, these included positive regulation of epithelial-to-mesenchymal transition (5 of 47 genes: *SDCBP*, *WWTR1*, *JAG1*, *MDK*, *TGFBR2*; adjusted p = 0.027), regulation of cell population proliferation, regulation of proteolysis and regulation of blood coagulation, together with the KEGG pathways tight junction and regulation of actin cytoskeleton (adjusted p = 0.034 each). The KEGG disease pathways in this list (Parkinson, Alzheimer and Huntington disease, amyotrophic lateral sclerosis, prion disease) were driven by shared proteasome, tubulin and oxidative-phosphorylation genes and do not indicate neurodegeneration-specific biology.

**Table 8.** Significant Enrichr terms (adjusted p < 0.05)

| Term | Overlap | Adjusted p |
| --- | --- | --- |
| **Top-200 DE genes — GO BP** |  |  |
| Macromolecule Biosynthetic Process (GO:0009059) | 12/183 | $2.85 \times 10^{-4}$ |
| Cytoplasmic Translation (GO:0002181) | 9/93 | $2.85 \times 10^{-4}$ |
| Translation (GO:0006412) | 12/234 | 0.002 |
| Peptide Biosynthetic Process (GO:0043043) | 10/158 | 0.002 |
| Positive Regulation Of Epithelial To Mesenchymal Transition (GO:0010718) | 5/47 | 0.027 |
| Positive Regulation Of Cell Population Proliferation (GO:0008284) | 15/483 | 0.027 |
| Regulation Of Proteolysis (GO:0030162) | 5/52 | 0.030 |
| Regulation Of Epithelial To Mesenchymal Transition (GO:0010717) | 6/83 | 0.030 |
| Regulation Of Blood Coagulation (GO:0030193) | 4/29 | 0.030 |
| Gene Expression (GO:0010467) | 11/296 | 0.030 |
| **Top-200 DE genes — KEGG** |  |  |
| Parkinson disease | 12/249 | 0.002 |
| Alzheimer disease | 14/369 | 0.002 |
| Coronavirus disease | 11/232 | 0.002 |
| Ribosome | 9/158 | 0.002 |
| Huntington disease | 11/306 | 0.010 |
| Amyotrophic lateral sclerosis | 12/364 | 0.010 |
| Pathways of neurodegeneration | 14/475 | 0.010 |
| Prion disease | 10/273 | 0.013 |
| Salmonella infection | 9/249 | 0.024 |
| Tight junction | 7/169 | 0.034 |
| Regulation of actin cytoskeleton | 8/218 | 0.034 |
| **Top-50 SHAP genes — KEGG** |  |  |
| Ribosome | 26/158 | $3.13 \times 10^{-40}$ |
| Coronavirus disease | 27/232 | $5.96 \times 10^{-38}$ |
| Oxidative phosphorylation | 6/133 | $3.86 \times 10^{-5}$ |
| Thermogenesis | 7/232 | $5.06 \times 10^{-5}$ |
| Parkinson disease | 7/249 | $6.47 \times 10^{-5}$ |
| Huntington disease | 7/306 | $2.08 \times 10^{-4}$ |

Sources: GO\_Enrichment\_TNBC.csv, KEGG\_TNBC.csv (top-200 DE genes) and KEGG\_Enrichment\_Results.csv (top-50 SHAP genes; first six of twelve significant pathways shown). Overlap = input genes in the term / term size. Enrichr default background.

The 50 top SHAP genes were enriched for 29 GO terms and 12 KEGG pathways, led by cytoplasmic translation (26 of 93 genes; adjusted p = 1.6×10^−46^) and the KEGG ribosome pathway (26 of 158; adjusted p = 3.1×10^−40^), followed by oxidative phosphorylation (Figure 8). This enrichment restates the composition of the input list, which was dominated by ribosomal and mitochondrial genes. For the 20 top High-Invasive lineage markers, no GO or KEGG term reached an adjusted p-value below 0.05 (smallest adjusted p = 0.16).

**Figure 8.**
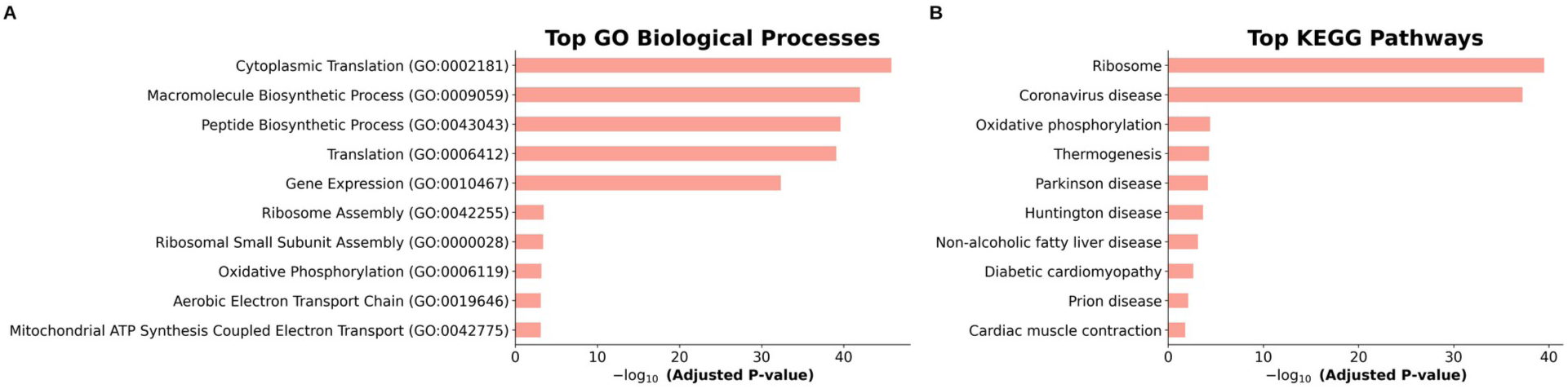
Enrichment of the 50 genes with the largest mean absolute SHAP values. (A) Top GO Biological Process terms. (B) Top KEGG pathways. Bars show −log_10_ of the Enrichr adjusted p-value.

### Rank-based enrichment of the sample-level comparison

GSEA of the full sample-level ranking identified nine Hallmark gene sets at FDR < 0.25 (Figure 9). KRAS signalling up (NES 2.30, FDR 0.001), E2F targets (NES 1.62, FDR 0.038), TNF-α signalling via NF-κB (NES 1.62, FDR 0.053), coagulation (NES 1.46, FDR 0.079) and the G2-M checkpoint (NES 1.30, FDR 0.18) were higher in invasive samples, whereas interferon-α response (NES −1.97, FDR 0.001), interferon-γ response (NES −1.65, FDR 0.019), myogenesis and apical surface were lower (Supplementary Table S2). The leading edge of the KRAS set included *PLAUR*, *ITGA2*, *EREG*, *HBEGF*, *DUSP6* and *ETV1*, and that of the E2F set included *HMGA1*, *MKI67*, *CDK1* and *TOP2A*. Among GO Biological Process terms, Hippo signalling was lower (NES −2.01, FDR 0.029) and positive regulation of lamellipodium assembly higher (NES 2.08, FDR 0.053) in invasive samples; no KEGG pathway reached FDR < 0.25. Over-representation analysis of all 6,651 sample-level significant genes against the tested background found no significant term (smallest adjusted p = 0.33), as expected for a list containing more than half of the tested genes.

**Figure 9.**
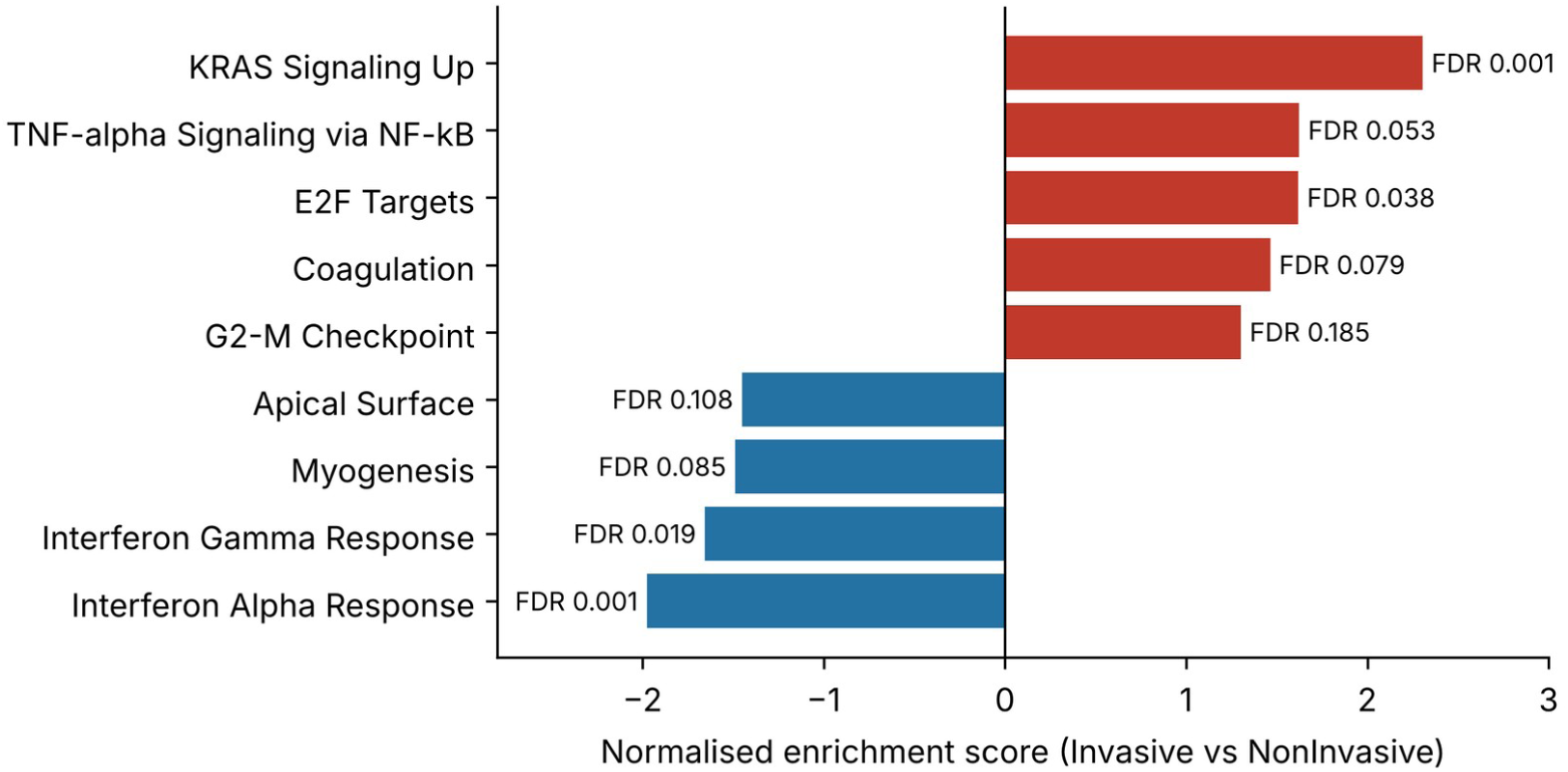
Hallmark gene sets with FDR < 0.25 in GSEA of the sample-level t statistic (invasive versus non-invasive samples). Positive NES, higher in invasive samples.

### High-Invasive lineage markers

Comparing the High-Invasive lineage group with all other cells ranked *ANXA5*, *SLC9A3R2*, *S100A6*, *TNFRSF12A*, *ENO1*, *C19orf33*, *S100A16*, *PRSS3* highest (Table 9; Figure 10). Most fold changes were small (0.03–0.59), with larger values for *PRSS3* (1.08) and *EPHB6* (2.63). This list overlapped with the 200 top-ranked invasive-versus-non-invasive genes in four genes (*METTL9*, *PRSS3*, *S100A16*, *SH3BGRL3*) and with the 50 top SHAP genes in none. In an enrichment analysis of the 100 top-ranked High-Invasive genes (notebook output; result file not supplied), the KEGG pathways bacterial invasion of epithelial cells (*ITGB1*, *CDC42*, *ARPC1B*, *ARPC1A*, *MET*; adjusted p = 0.0054), ribosome (0.0092) and ubiquitin-mediated proteolysis (0.030) were significant. Because the comparison group contains all other cells, including parental cells, these markers describe the High-Invasive lineage relative to a mixed background rather than relative to non-invasive cells.

**Figure 10.**
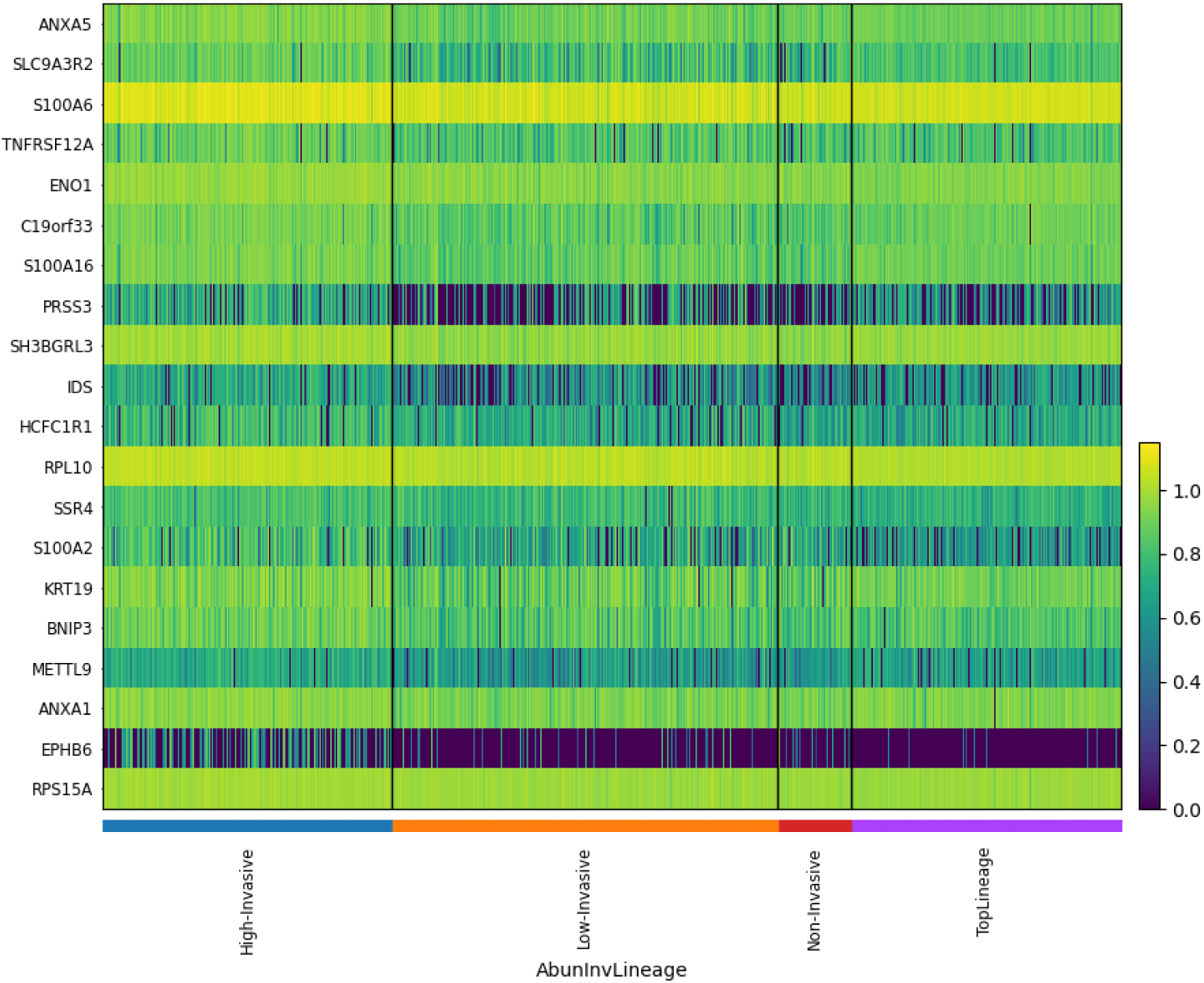
Expression of the 20 top High-Invasive markers across lineage groups (data generators’ annotation).

**Table 9.** Top 20 marker genes of the High-Invasive lineage group (versus all other cells)

| Gene | Score | Log fold change | Gene | Score | Log fold change |
| --- | --- | --- | --- | --- | --- |
| *ANXA5* | 62.7 | 0.127 | *HCFC1R1* | 46.5 | 0.274 |
| *SLC9A3R2* | 55.8 | 0.261 | *RPL10* | 46.3 | 0.044 |
| *S100A6* | 55.2 | 0.062 | *SSR4* | 46.1 | 0.176 |
| *TNFRSF12A* | 52.6 | 0.217 | *S100A2* | 44.7 | 0.429 |
| *ENO1* | 51.9 | 0.076 | *KRT19* | 44.2 | 0.173 |
| *C19orf33* | 51.2 | 0.145 | *BNIP3* | 44.1 | 0.170 |
| *S100A16* | 51.1 | 0.130 | *METTL9* | 44.0 | 0.324 |
| *PRSS3* | 49.1 | 1.082 | *ANXA1* | 43.8 | 0.080 |
| *SH3BGRL3* | 48.2 | 0.070 | *EPHB6* | 43.4 | 2.631 |
| *IDS* | 46.6 | 0.594 | *RPS15A* | 43.0 | 0.031 |

Source: Top20\_HighInvasive\_Biomarkers.csv; all adjusted p-values were reported as 0 (cell-level Wilcoxon test).

### Protein interaction network

The ten genes with the highest node degree in the STRING network were all ribosomal proteins (*RPL18A*, degree 33; *RPL15*, *RPL24*, *RPL35* and *RPS3*, 32; *RPL3*, *RPL36*, *RPL36A*, *RPL38* and *RPL39*, 31; Supplementary Figure S3). Eight of these genes are among the 50 top SHAP genes, but *RPL24* and *RPL36A* are not in any result list supplied, so the network input cannot be reconstructed. Ribosomal proteins form a large, densely connected complex in STRING, so high degree is expected for them and does not by itself indicate a role in invasion.

### Comparison with published evidence

Table 10 summarises published evidence for selected candidates. Experimental studies link *HMGA1* to epithelial-to-mesenchymal transition, migration, invasion and metastasis of basal-like breast cancer cells [23], and *ANXA1* to TGF-β signalling and metastasis of basal-like cells [24]. *VIM* is an established mesenchymal marker [25]. For *EPHB6*, the evidence points in more than one direction: in TNBC models EPHB6 accelerated tumour growth and initiation but partially suppressed epithelial-to-mesenchymal transition and increased sensitivity to DNA-damaging therapy [26]. Loss rather than gain of *BNIP3* promoted mammary tumour progression to metastasis in a mouse model [27], whereas *BNIP3* was modestly higher in the High-Invasive group here and did not differ between invasive and non-invasive samples (FDR 0.14). Post-translational regulation of *ENO1* has been described in TNBC cells [28], but ENO1 did not differ at the sample level (FDR 0.76). For *ANXA5* and *KRT19* we did not find a study that tested their expression as a TNBC invasion marker; the earlier interpretation table, which marked all five of ANXA5, ENO1, BNIP3, KRT19 and EPHB6 as ‘reported in TNBC’, did not cite sources and should not be used.

**Table 10.** Published evidence for selected candidate genes.

| Gene | Evidence in this study | Published finding | Evidence type | Ref. |
| --- | --- | --- | --- | --- |
| *VIM* | Higher in invasive samples (FDR 0.001) | Mesenchymal intermediate filament; marker of EMT and motility | Review | [25] |
| *HMGA1* | Higher in invasive samples (FDR 0.002); E2F leading edge | Knock-down reverses EMT and reduces migration, invasion and metastasis in basal-like cells | Experimental | [23] |
| *ANXA1* | High-Invasive marker; higher in invasive samples (FDR 0.011) | Promotes TGF-β signalling and metastasis of basal-like breast cancer cells | Experimental | [24] |
| *EPHB6* | High-Invasive marker; held-out SHAP rank 16; sample-level FDR 0.31 | Promotes TNBC tumour growth and initiation; partially suppresses EMT; increases drug sensitivity | Experimental | [26] |
| *BNIP3* | High-Invasive marker; sample-level FDR 0.14 | Loss promotes mammary tumour progression to metastasis (direction opposite to this study) | Experimental (mouse) | [27] |
| *ENO1* | High-Invasive marker; sample-level FDR 0.76 | Ser419 phosphorylation drives nuclear export in TNBC cells | Experimental | [28] |
| *ANXA5* | High-Invasive marker (rank 1) | No study of expression as a TNBC invasion marker found | — | — |
| *KRT19* | High-Invasive marker (rank 15) | No TNBC-specific invasion study found | — | — |

Only studies with verified bibliographic details are cited. Computational prioritisation in this study is not independent validation; agreement with the literature indicates biological plausibility only.

## Discussion

We re-analysed single-cell transcriptomes of MDA-MB-231 cells separated by a transwell invasion assay, first at the level of individual cells and then with every test and model evaluated on independent samples. The sample-aware analysis changed the interpretation in two ways. It confirmed that invasive and non-invasive cells differ reproducibly: the main cell-level genes remained significant across the five pairs of samples, and a classifier trained on some samples recognised the invasive state in samples it had never seen (AUC 0.92). It also showed that the ribosomal and mitochondrial genes that dominated the original SHAP ranking were not needed for this separation; once leakage was removed and interpretation was restricted to held-out samples, the model relied on inflammatory, stress-response and EMT-related genes.

The biology of the invasive state is plausible. Genes higher in invasive samples included *VIM* [25] and *HMGA1*, which promotes EMT, invasion and metastasis in basal-like breast cancer cells [23], together with the EMT regulators *TGFBR2*, *JAG1* and *WWTR1*. Rank-based enrichment pointed to KRAS-associated signalling, with leading-edge genes such as *PLAUR*, *ITGA2* and *HBEGF* that are linked to adhesion, matrix remodelling and growth-factor signalling, and to E2F-driven proliferation genes, while Hippo signalling and interferon responses were lower. MDA-MB-231 carries an activating *KRAS* mutation, so differences in KRAS-pathway output between subpopulations are consistent with the cell line’s biology, although our data cannot show whether these programmes cause invasion or follow from it.

Two alternative explanations must be considered. First, all invasive samples were sequenced in one library and all non-invasive samples in another, so the comparison is confounded with library. Holding out whole samples removes leakage between samples but not this confounding. The weak performance of the QC-metric model (AUC 0.63) argues that the main measured technical differences explain little of the separation, but unmeasured library effects cannot be excluded. Second, invasive cells were recovered from the underside of the membrane and non-invasive cells from the upper chamber, so differences in handling could induce immediate-early and stress-response genes such as *JUN*, *DUSP1* and *DDIT4*. These genes ranked highly in the held-out SHAP analysis, and their contribution should be treated with caution. SHAP values describe what a model uses [10,11]; they depend on the chosen fold and features and do not establish causal roles.

The lineage annotation adds a clonal dimension. The High-Invasive lineage markers overlapped little with the invasive-versus-non-invasive genes, and several, including *ENO1* and *BNIP3*, did not differ between invasive and non-invasive samples, which suggests that clone-level invasiveness and the population-level invasive state capture different signals. *EPHB6* was both a High-Invasive marker and a held-out SHAP feature; published work in TNBC models links it to tumour growth and initiation but also to partial suppression of EMT [26], so its role in invasion remains open. The earlier mechanism diagram, which connected selected markers to HIF-1 signalling, glycolysis, autophagy and mitophagy, was based on single-gene, non-significant pathway terms and was not retained.

Methodologically, this study illustrates how the conclusions of an explainable-AI analysis depend on the validation design. The original random cell split produced an accuracy of 96.6% and a SHAP ranking dominated by ribosomal genes; grouped validation produced a lower but more credible AUC and a biologically more coherent set of features. Reporting sample-level statistics, grouped validation and simple technical baselines alongside SHAP is therefore essential [12–14].

### Limitations and Future Work

This study has several limitations. First, all data come from one TNBC cell line in one in vitro assay, so the findings describe invasive and non-invasive MDA-MB-231 cells and cannot be generalised to TNBC patients without validation. Second, the invasive, non-invasive and parental populations were each sequenced in a separate library, so biological and library effects cannot be fully separated, and the parental population consists of a single sample. Third, cells from the two chambers were harvested from different surfaces, which may induce stress-response genes. Fourth, only compressed normalised values were available and no raw counts, so count-based methods could not be used and effect sizes are reported on the deposited scale. Fifth, replication is limited to five samples per group from one experiment, and samples sharing a well number may not be fully independent. Sixth, held-out SHAP values were computed for one fold, and rankings may vary between folds. Seventh, the original SHAP cells were sampled without a fixed seed and the STRING analysis was not documented. Finally, no independent dataset or experimental validation was performed.

Future work should repeat the transwell experiment with sample-level library multiplexing, so that invasive and non-invasive cells are sequenced together; test the candidate genes in other TNBC cell lines and in patient single-cell datasets; and perturb selected genes, such as *HMGA1*, *TGFBR2* and the KRAS-associated leading-edge genes, to establish whether they influence invasion.

## CONCLUSION

Invasive and non-invasive MDA-MB-231 cells from a transwell assay differ in a transcriptional state that is reproducible across independent samples and can be recognised by an XGBoost classifier in samples withheld from training (AUC 0.92). The state involves EMT-related genes such as *VIM*, *HMGA1* and *TGFBR2*, higher KRAS-associated and E2F-driven programmes, and lower interferon responses. Because the populations were sequenced in separate libraries and harvested differently, and only one cell line was studied, these genes are candidates rather than validated biomarkers. Independent replicate experiments, other TNBC cell lines and patient datasets, and functional perturbation are needed to establish their role in invasion.

## DECLARATIONS

### Data availability

The single-cell RNA-seq data analysed in this study are publicly available from the NCBI Gene Expression Omnibus under accession GSE329526 (https://www.ncbi.nlm.nih.gov/geo/query/acc.cgi?acc=GSE329526; processed file GSE329526_MB231.h5ad). Raw sequencing data are available from the NCBI Sequence Read Archive under BioProject PRJNA1459420 (https://www.ncbi.nlm.nih.gov/bioproject/PRJNA1459420).

### Code availability

The original analysis notebook, the sample-aware re-analysis code, all result tables and the supplementary data are openly available on GitHub at https://github.com/tanjidnijum02/tnbc-invasion-scrna-xai and archived on Zenodo (release v1.0.0; https://doi.org/10.5281/zenodo.23494607) under the MIT licence.

### Author contributions

T.H. and S.B. conceived the study, performed the analyses, interpreted the results and wrote the manuscript. Both authors read and approved the final manuscript.

### Ethics statement

This study used only publicly available data from a commercially available human cell line and did not involve human participants or animals; ethical approval was not required.

## Acknowledgements.

We thank the data generators (De Santiago C, Gardner AL, Parker P, Thomas JR, Callahan G, Ghosh M, Baker AB and Brock A) for making GSE329526 publicly available.

## SUPPLEMENTARY MATERIAL

The following supplementary material is provided in a separate file (Supplementary_Material_Huda_Biswas.pdf/.docx) and as a spreadsheet (Supplementary_Tables_Huda_Biswas.xlsx):

- **Supplementary Figure S1.** UMAP embedding coloured by Leiden cluster.
- **Supplementary Figure S2.** Mean expression of the 20 top SHAP genes of the original model per label.
- **Supplementary Figure S3.** Node degree of the ten top-ranked genes in the STRING network.
- **Supplementary Table S1.** Per-sample predictions of the three grouped cross-validation models.
- **Supplementary Table S2.** Hallmark GSEA results for all gene sets.
- **Supplementary Table S3.** Top 50 genes by mean absolute SHAP value on held-out samples.
- **Supplementary Table S4.** Sample-level differential expression for all 11,727 tested genes (spreadsheet only).

