## Supplementary Material (Figures S1-S3 and Tables S1-S3) for "Integrative Single-Cell Transcriptomics and Explainable AI for Disease Mechanism and Biomarker Discovery in Triple-Negative Breast Cancer"

### SUPPLEMENTARY FIGURES

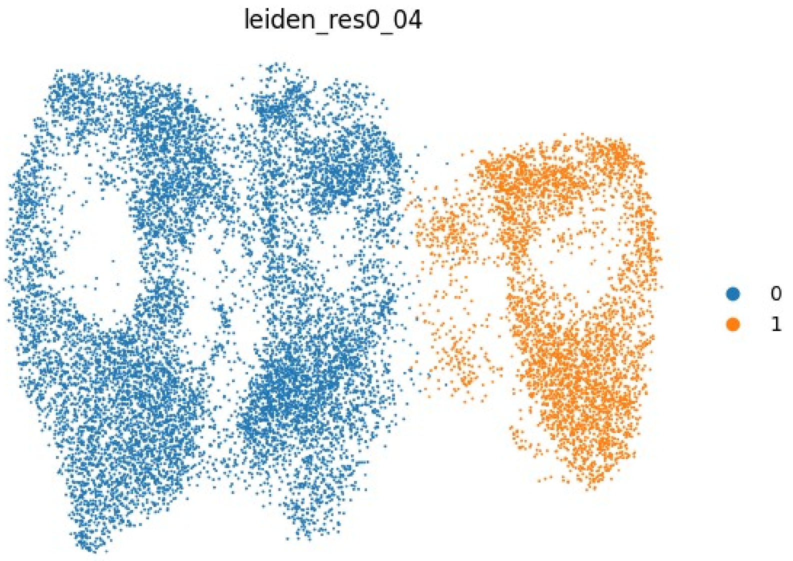


**Supplementary Figure S1. UMAP embedding provided in the deposited object (GSE329526_MB231.h5ad), coloured by Leiden cluster (resolution 0.04; data generators' annotation).**

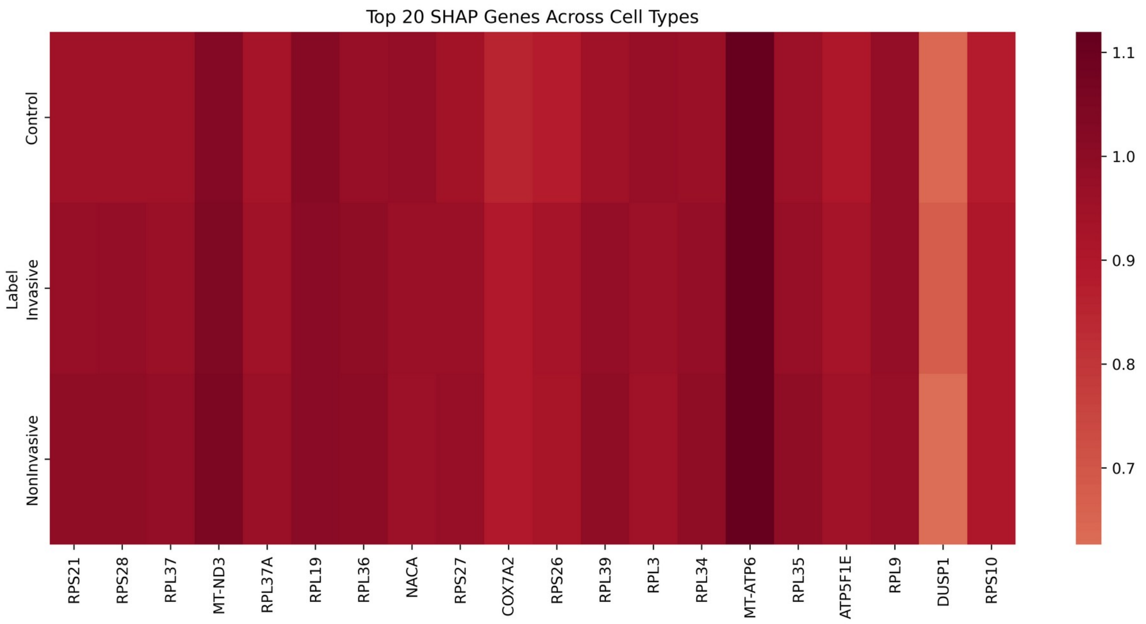


**Supplementary Figure S2. Mean expression of the 20 top SHAP genes of the original three-class model per label (Control = parental, Invasive, NonInvasive). Differences between labels are a few percent and are barely visible on this colour scale.**

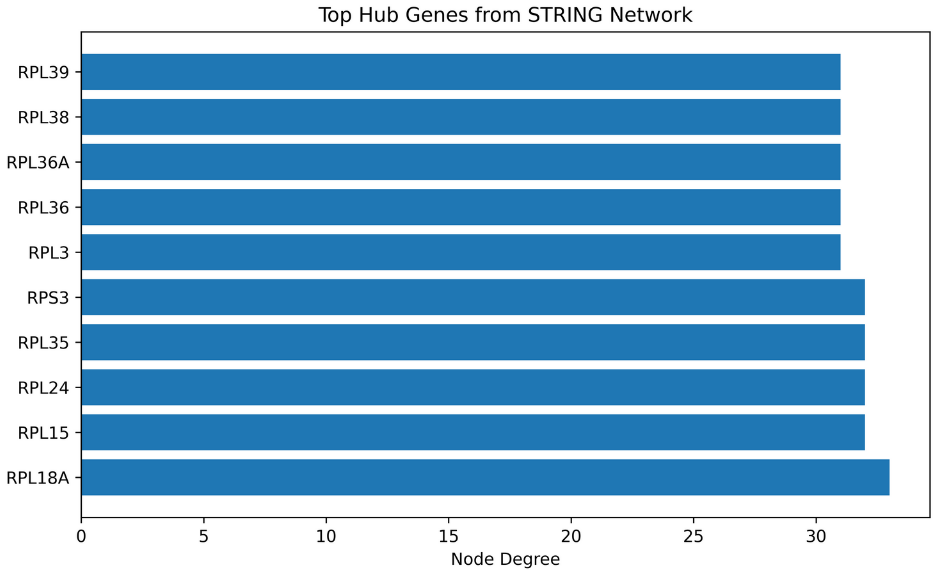


**Supplementary Figure S3. Node degree of the ten top-ranked genes in the STRING network. Degree values were entered manually from STRING; the input gene list and network settings were not recorded, so this result is descriptive only.**

### SUPPLEMENTARY TABLES

#### Supplementary Table S1. Predictions for every held-out sample in sample-grouped five-fold cross-validation

| Model | Held-out sample | Label | Mean P(invasive) | Cells called invasive (%) |
|---|---|---|---|---|
| HVGs | B3 | NonInvasive | 0.275 | 21.2 |
| HVGs | B5 | NonInvasive | 0.216 | 14.4 |
| HVGs | B6 | NonInvasive | 0.201 | 12.5 |
| HVGs | B7 | NonInvasive | 0.243 | 18.2 |
| HVGs | B8 | NonInvasive | 0.236 | 16.6 |
| HVGs | C3 | Invasive | 0.758 | 82.6 |
| HVGs | C5 | Invasive | 0.786 | 88.7 |
| HVGs | C6 | Invasive | 0.759 | 82.2 |
| HVGs | C7 | Invasive | 0.739 | 82.2 |
| HVGs | C8 | Invasive | 0.785 | 86.1 |
| HVGs without RP/MT | B3 | NonInvasive | 0.280 | 23.1 |
| HVGs without RP/MT | B5 | NonInvasive | 0.220 | 14.4 |
| HVGs without RP/MT | B6 | NonInvasive | 0.201 | 12.0 |
| HVGs without RP/MT | B7 | NonInvasive | 0.233 | 16.9 |
| HVGs without RP/MT | B8 | NonInvasive | 0.231 | 16.3 |
| HVGs without RP/MT | C3 | Invasive | 0.754 | 81.8 |
| HVGs without RP/MT | C5 | Invasive | 0.788 | 87.8 |
| HVGs without RP/MT | C6 | Invasive | 0.758 | 82.8 |
| HVGs without RP/MT | C7 | Invasive | 0.732 | 81.7 |
| HVGs without RP/MT | C8 | Invasive | 0.784 | 86.5 |
| QC metrics only | B3 | NonInvasive | 0.470 | 41.4 |
| QC metrics only | B5 | NonInvasive | 0.492 | 45.6 |
| QC metrics only | B6 | NonInvasive | 0.482 | 42.7 |
| QC metrics only | B7 | NonInvasive | 0.478 | 43.6 |
| QC metrics only | B8 | NonInvasive | 0.435 | 30.9 |
| QC metrics only | C3 | Invasive | 0.532 | 55.7 |
| QC metrics only | C5 | Invasive | 0.581 | 66.7 |
| QC metrics only | C6 | Invasive | 0.552 | 62.8 |
| QC metrics only | C7 | Invasive | 0.555 | 61.5 |
| QC metrics only | C8 | Invasive | 0.522 | 52.3 |

Each fold held out one invasive and one non-invasive sample with the same well number. Source: xgb_*_per_sample.csv.

#### Supplementary Table S2. GSEA prerank of the sample-level t statistic against MSigDB Hallmark 2020 (all tested gene sets)

| Hallmark gene set | NES | Nominal p | FDR | Leading edge (tag %) |
|---|---|---|---|---|
| KRAS Signaling Up | +2.30 | 0.001 | 0.001 | 40/113 |
| Interferon Alpha Response | -1.97 | 0.001 | 0.001 | 35/83 |
| Interferon Gamma Response | -1.65 | 0.001 | 0.019 | 61/150 |
| E2F Targets | +1.62 | 0.001 | 0.038 | 79/197 |
| TNF-alpha Signaling via NF-kB | +1.62 | 0.001 | 0.053 | 36/167 |
| Coagulation | +1.46 | 0.001 | 0.079 | 16/66 |
| Myogenesis | -1.49 | 0.005 | 0.085 | 37/101 |
| Apical Surface | -1.45 | 0.048 | 0.108 | 11/26 |
| G2-M Checkpoint | +1.30 | 0.001 | 0.185 | 79/190 |
| Myc Targets V2 | -1.35 | 0.078 | 0.260 | 38/56 |

| Hallmark gene set | NES | Nominal p | FDR | Leading edge (tag %) |
|---|---|---|---|---|
| Wnt-beta Catenin Signaling | -1.29 | 0.120 | 0.418 | 13/30 |
| Notch Signaling | -1.23 | 0.177 | 0.502 | 8/25 |
| IL-6/JAK/STAT3 Signaling | -1.24 | 0.162 | 0.526 | 20/53 |
| Bile Acid Metabolism | -1.20 | 0.191 | 0.597 | 17/64 |
| Xenobiotic Metabolism | +1.04 | 0.375 | 0.644 | 16/124 |
| Estrogen Response Late | +1.00 | 0.667 | 0.686 | 37/137 |
| Apoptosis | +1.05 | 0.417 | 0.701 | 27/125 |
| Myc Targets V1 | +0.87 | 1.000 | 0.894 | 29/195 |
| PI3K/AKT/mTOR Signaling | +0.90 | 0.611 | 0.907 | 11/86 |
| TGF-beta Signaling | +0.80 | 0.889 | 0.946 | 13/49 |
| Spermatogenesis | +0.67 | 1.000 | 0.969 | 13/60 |
| Reactive Oxygen Species Pathway | -0.54 | 0.983 | 0.994 | 17/45 |
| Apical Junction | -0.97 | 0.563 | 1.000 | 38/130 |
| Androgen Response | -0.61 | 0.982 | 1.000 | 27/84 |
| Hedgehog Signaling | -1.11 | 0.336 | 1.000 | 10/20 |
| Pperoxisome | -0.69 | 0.926 | 1.000 | 16/80 |
| DNA Repair | -0.69 | 0.962 | 1.000 | 39/145 |
| KRAS Signaling Dn | -0.71 | 0.912 | 1.000 | 7/53 |
| Protein Secretion | -0.73 | 0.932 | 1.000 | 28/94 |
| Unfolded Protein Response | -0.74 | 0.911 | 1.000 | 40/108 |
| Angiogenesis | -0.75 | 0.809 | 1.000 | 4/17 |
| Mitotic Spindle | -0.75 | 0.930 | 1.000 | 42/193 |
| Complement | -0.76 | 0.923 | 1.000 | 39/127 |
| heme Metabolism | -0.77 | 0.890 | 1.000 | 41/145 |
| Epithelial Mesenchymal Transition | -1.00 | 0.498 | 1.000 | 31/128 |
| Hypoxia | -1.08 | 0.292 | 1.000 | 51/162 |
| UV Response Up | -0.82 | 0.838 | 1.000 | 43/122 |
| Oxidative Phosphorylation | -0.86 | 0.797 | 1.000 | 54/198 |
| Adipogenesis | -1.06 | 0.344 | 1.000 | 67/170 |
| Glycolysis | -0.88 | 0.791 | 1.000 | 52/167 |
| UV Response Dn | -0.88 | 0.746 | 1.000 | 36/122 |
| Inflammatory Response | -0.89 | 0.722 | 1.000 | 27/108 |
| Allograft Rejection | -1.03 | 0.454 | 1.000 | 42/102 |
| Fatty Acid Metabolism | -0.91 | 0.683 | 1.000 | 24/122 |
| IL-2/STAT5 Signaling | -0.92 | 0.669 | 1.000 | 38/142 |
| mTORC1 Signaling | -0.61 | 0.994 | 1.000 | 45/190 |
| Cholesterol Homeostasis | -0.95 | 0.614 | 1.000 | 13/66 |
| p53 Pathway | -0.81 | 0.864 | 1.000 | 31/166 |
| Estrogen Response Early | -0.94 | 0.627 | 1.000 | 39/142 |

Positive NES, higher in invasive samples. 1,000 permutations, seed 42. Source: gsea_prerank_MSigDB_Hallmark_2020.csv.

### Supplementary Table S3. Top 50 genes by mean absolute SHAP value on held-out samples B7 and C7

| Rank | Gene | Mean \|SHAP\| | Sample-level difference | FDR |
|---|---|---|---|---|
| 1 | *CCL2* | 0.803 | -0.064 | 0.00046 |
| 2 | *LGALS12* | 0.576 | -0.205 | 0.00088 |
| 3 | *DUSP1* | 0.423 | +0.054 | 0.034 |
| 4 | *MTRNR2L8* | 0.240 | -0.143 | 0.00014 |
| 5 | *PRSS2* | 0.237 | -0.105 | 0.016 |
| 6 | *CXCL3* | 0.215 | +0.127 | 0.004 |
| 7 | *KYNU* | 0.214 | +0.170 | 0.0019 |
| 8 | *AL603756.1* | 0.207 | +0.088 | 0.001 |
| 9 | *CXCL2* | 0.189 | +0.080 | 0.03 |
| 10 | *DDIT4* | 0.184 | +0.014 | 0.027 |
| 11 | *ATP2B1-AS1* | 0.156 | +0.069 | 0.038 |
| 12 | *NFKBIZ* | 0.151 | +0.123 | 0.0014 |
| 13 | *LSR* | 0.150 | -0.141 | 0.0012 |
| 14 | *POU2F2* | 0.145 | +0.090 | 0.0022 |
| 15 | *JUN* | 0.136 | +0.043 | 0.025 |
| 16 | *EPHB6* | 0.126 | +0.035 | 0.31 |
| 17 | *NEDD9* | 0.124 | -0.222 | 0.00088 |
| 18 | *AGR2* | 0.121 | +0.206 | 0.0025 |
| 19 | *CCDC3* | 0.120 | -0.043 | 0.00045 |
| 20 | *IFITM3* | 0.120 | -0.033 | 0.0052 |
| 21 | *AC007952.4* | 0.115 | +0.050 | 0.024 |
| 22 | *DUSP6* | 0.112 | +0.045 | 0.0018 |
| 23 | *CXCL8* | 0.106 | +0.085 | 0.017 |
| 24 | *IFITM1* | 0.105 | -0.019 | 0.0042 |
| 25 | *SPINK4* | 0.104 | -0.084 | 0.0041 |
| 26 | *IL24* | 0.095 | -0.103 | 0.0013 |
| 27 | *MAP3K8* | 0.092 | -0.001 | 0.94 |
| 28 | *AC016831.7* | 0.091 | -0.010 | 0.088 |
| 29 | *MT-ND4L* | 0.091 | -0.033 | 0.0011 |
| 30 | *EPAS1* | 0.091 | -0.153 | 0.0026 |
| 31 | *MANCR* | 0.089 | +0.190 | 0.0022 |
| 32 | *HIST1H4C* | 0.088 | -0.025 | 0.036 |
| 33 | *TNFSF10* | 0.087 | -0.241 | 0.0013 |
| 34 | *C15orf48* | 0.085 | -0.021 | 0.22 |
| 35 | *HSPA8* | 0.085 | -0.008 | 0.36 |
| 36 | *ZG16B* | 0.081 | +0.163 | 0.00088 |
| 37 | *C3* | 0.080 | -0.078 | 0.0053 |
| 38 | *F3* | 0.080 | -0.141 | 0.0025 |
| 39 | *HMGA2-AS1* | 0.079 | +0.009 | 0.045 |
| 40 | *ADAM8* | 0.078 | +0.086 | 0.0046 |
| 41 | *GFPT2* | 0.077 | +0.037 | 0.0021 |
| 42 | *CSF3* | 0.077 | — | — |
| 43 | *MT-ATP8* | 0.076 | -0.015 | 0.12 |
| 44 | *OTULINL* | 0.072 | +0.110 | 0.0023 |
| 45 | *FSIP2* | 0.072 | +0.122 | 0.0029 |
| 46 | *ASB2* | 0.072 | — | — |
| 47 | *XAF1* | 0.067 | -0.156 | 0.002 |
| 48 | *TGFBI* | 0.067 | -0.121 | 0.0026 |
| 49 | *AC020916.1* | 0.065 | +0.027 | 0.25 |
| 50 | *CCNL1* | 0.064 | +0.032 | 0.0082 |

1,000 cells, seed 42. Difference, invasive minus non-invasive sample mean on the deposited normalised scale; — , gene not in the sample-level test (detected in <10% of cells).

**Supplementary Table S4.** Sample-level differential expression (five invasive versus five non-invasive samples; Welch t-test, Benjamini–Hochberg FDR) for all 11,727 tested genes is provided in the spreadsheet Supplementary_Tables_Huda_Biswas.xlsx (sheet S4).
